# Mechanisms underlying sequence-dependent DNA hybridisation rates in the absence of secondary structure

**DOI:** 10.1101/2021.12.17.473246

**Authors:** Sophie Hertel, Richard E. Spinney, Stephanie Y. Xu, Thomas E. Ouldridge, Richard G. Morris, Lawrence K. Lee

**Affiliations:** EMBL Australia Node for Single Molecule Science, School of Medical Sciences, UNSW Sydney, 2052, Australia; School of Physics, University of New South Wales - Sydney 2052, Australia; Department of Bioengineering and Centre for Synthetic Biology, Imperial College London, London, SW7 2AZ, United Kingdom; ARC Centre of Excellence in Synthetic Biology, University of New South Wales, Sydney, Australia

## Abstract

The kinetics of DNA hybridisation are fundamental to biological processes and DNA-based technologies. However, the precise physical mechanisms that determine why different DNA sequences hybridise at different rates are not well understood. Secondary structure is one predictable factor that influences hybridisation rates but is not sufficient on its own to fully explain the observed sequence-dependent variance. Consequently, to achieve a good correlation with experimental data, current prediction algorithms require many parameters that provide little mechanistic insight into DNA hybridisation. In this context, we measured hybridisation rates of 43 different DNA sequences that are not predicted to form secondary structure and present a parsimonious physically justified model to quantify their hybridisation rates. Accounting only for the combinatorics of complementary nucleating interactions and their sequence-dependent stability, the model achieves good correlation with experiment with only two free parameters, thus providing new insight into the physical factors underpinning DNA hybridisation rates.

## INTRODUCTION

DNA is a biopolymer formed from four different nucleotides, adenine, thymine, guanine and cytosine (A,T,G and C respectively), whose order or sequence is used to encode information that is the foundation of biology. Complementary DNA strands hybridise via Watson and Crick base pairing between A-T or G-C bases to form the DNA double helix or duplex (1), whose structural (2) and physical (3-5) properties are well characterised. In addition to its essential role in biology, DNA hybridisation also underpins DNA nanotechnology (6-8), which utilises DNA self-assembly for the construction of rationally designed nanoscale structures and machines (9-16). DNA nanotechnology has led to the development of a broad range of technologies including applications in molecular sensing (17-20), coordinating complex reaction cascades (21-23), drug delivery vessels (24-27) and super resolution imaging methods, such as DNA points accumulation for imaging in nanoscale topography (DNA-PAINT) (28). Thus, understanding the thermodynamics, kinetics and mechanisms for DNA hybridisation is fundamentally important for biology and biotechnology.

The thermodynamics of DNA hybridisation have long been observable via spectrophotometric or viscometric observations of thermal melt curves and are well studied (29-31). The reaction is dominated by states consisting of completely dissociated DNA strands or fully hybridised DNA duplexes. The stability of a DNA duplex can therefore be estimated from the structure of a fully hybridised duplex and is dependent on hydrogen bonds between paired bases in DNA duplexes and hydrophobic base stacking that occurs between neighbouring base pairs; both these interactions are sequence dependent (32, 33). Models predicting the melting temperature *T*_*m*_ of a DNA duplex adopt a two-state nearest neighbour approach, which postulates that the stability of a given base pair depends on the identity of the nucleotide bases involved (A-T or G-C) and its nearest neighbour base pairs. In turn, the *T*_*m*_ associated with the formation of any DNA duplex can be estimated from the sum of the free energy of all 2 contiguous base-pairing interactions, as well as additional parameters to account for the relative stabilities of the ends of the duplex (32, 33). Given that there are only 10 unique combinations of 2-base sequences, nearest neighbour models can be parameterised experimentally including in different buffer conditions (34-41), and algorithmic implementations predict hybridisation *T*_*m*_ reasonably well (42, 43).

The *kinetics* of DNA hybridisation are also sequence-dependent (44-47). However, the pathways to DNA hybridisation are difficult to observe, and therefore the physical basis for sequence-dependent hybridisation rates remains poorly understood. Based on early thermodynamic and kinetic measurements of DNA hybridisation, Pörschke and colleagues proposed a reaction mechanism in which hybridisation proceeds via a slow, rate limiting bimolecular nucleation step, followed by fast monomolecular ‘zippering’ into a fully formed DNA duplex (30, 48, 49). More recent developments in coarse-grained molecular dynamics (MD) simulations enabled an *in silico* view of hybridisation pathways, in which the rate-limiting nucleation step consisted of a short stretch (∼3 bp at 300 K) of contiguous and complementary base-pairing interactions (50-52). Since there are typically many such possible nucleating interactions, it therefore follows that the combination and relative stability of all possible nucleating interactions, which is entirely determined by the DNA sequence, defines the overall activation free energy and therefore the rate of any DNA hybridisation reaction. However, DNA strands can also form intramolecular interactions that result in secondary structures such as hairpins that influence both the rates of hybridisation and melting. Such secondary structure can reduce hybridisation rates either by limiting the availability of a subset of nucleating interactions or by lowering the probability that any given nucleating interaction is stable enough to favour the displacement of the secondary structure, which must be denatured prior to zippering into a fully formed duplex (53-55).

Two algorithms have recently been developed for predicting sequence dependent hybridisation rates. A ‘weighted neighbour voting’ algorithm was used to examine 50 different sequence-dependent physical ‘features’ and found 35 different features that correlated with hybridisation rates (56). Perhaps unsurprisingly, of these features, the ensemble standard free energy of secondary structure emerged as the single best predictor of DNA hybridisation rates, reporting predictions of hybridisation rate constants (*k*_*a*_) of ∼60% accuracy within a factor of two. The inclusion of five additional features resulted in a six-parameter model, which achieved a reported ∼80% prediction accuracy within a factor of two. However, apart from secondary structure, the physical mechanisms underpinning how these additional features influence hybridisation rates remain unclear. Hata *et al*. subsequently presented an alternative, physically motivated, model by estimating the relative binding capability for all 3 consecutive base sequences involved in all possible nucleation interactions, including those which were off-register or mis-matched (45). This capability was dependent on an estimate of the propensity of any of the 32 possible 3-base nucleating interactions to seed full hybridisation and on the probability of predicted secondary structures sterically hindering nucleating interactions.

Surprisingly however, seeding propensities did not correlate with the stability of nucleating interactions and accurate predictions required these propensities to be determined empirically by fitting 32 free parameters to experimental data. Thus, secondary structure remains the only physically well-defined determinant for algorithms predicting sequence dependent hybridisation rates. However, accurate predictions require multiple additional parameters that are not physically well defined. This suggests that there are other dominating physical factors apart from secondary structure that are yet to be identified.

Coarse grained MD simulations, for example, indicate that nucleation can occur from base pairing interactions that are off-register from a fully formed duplex (50-52). In these instances, off-register nucleation states can progress to metastable intermediaries such as misaligned duplexes that can move into register via inchworming or pseudoknot internal displacement mechanisms, followed by the final zippering step into the fully hybridized DNA double strand (50). Like zippering, these monomolecular rearrangements also occur much more rapidly than nucleation. Consequently, repetitive sequences, which have a greater number of possible off-register nucleating interactions were predicted to hybridise more rapidly than non-repetitive sequences (50).

Here we explored the impact of off-register nucleating states on the hybridisation rate of DNA strands experimentally. To reduce the complexity of hybridisation pathways and to identify physical elements yet to be explicitly accounted for in predictive models, we focused on sequences that were not predicted to form secondary structures. Using surface plasmon resonance (SPR) we measured the hybridisation rates of 43 different DNA strands with varying GC content and degree of sequence repetition and demonstrate that repetitive sequences do indeed hybridise more rapidly than non-repetitive sequences. We also present a simple, physically-justified model, which demonstrates that it is possible to capture much of the variance in sequence dependent hybridisation rates with only two free parameters that account for the combination and stability of all possible nucleating interactions, including those that are off-register.

## MATERIALS AND METHODS

### DNA oligonucleotides

All DNA was purchased from IDT. The salt purified oligonucleotides were resuspended in milliQwater and stored at -20 °C. To ensure that the measured hybridization kinetics only depended on differences in the sequence, DNA strands were designed to have no or negligible secondary structures (2bp or less) and nearly the same free energy of the lowest energy double stranded complex, using NUPACK and IDT (Table 1).

**Table 1.**
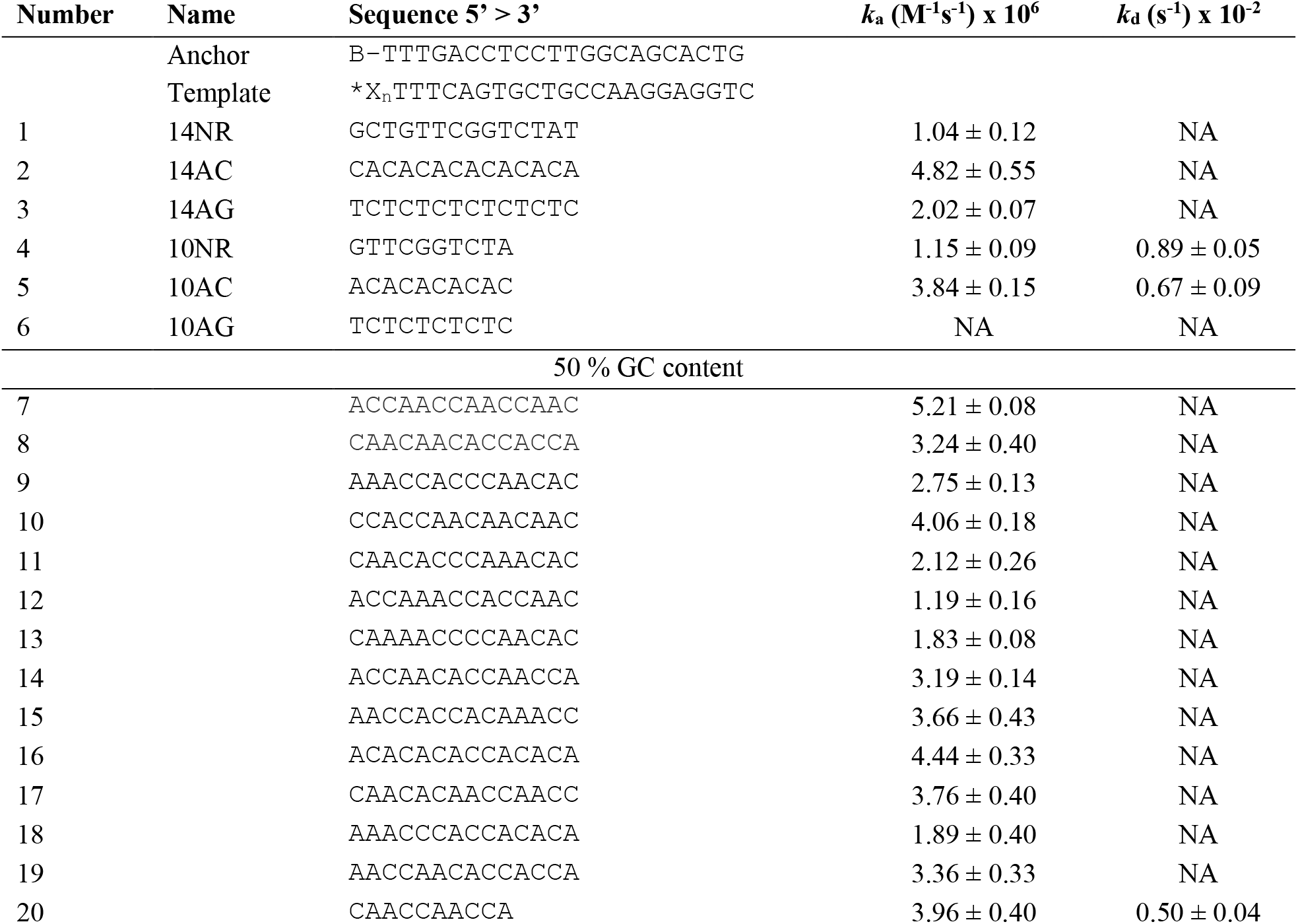
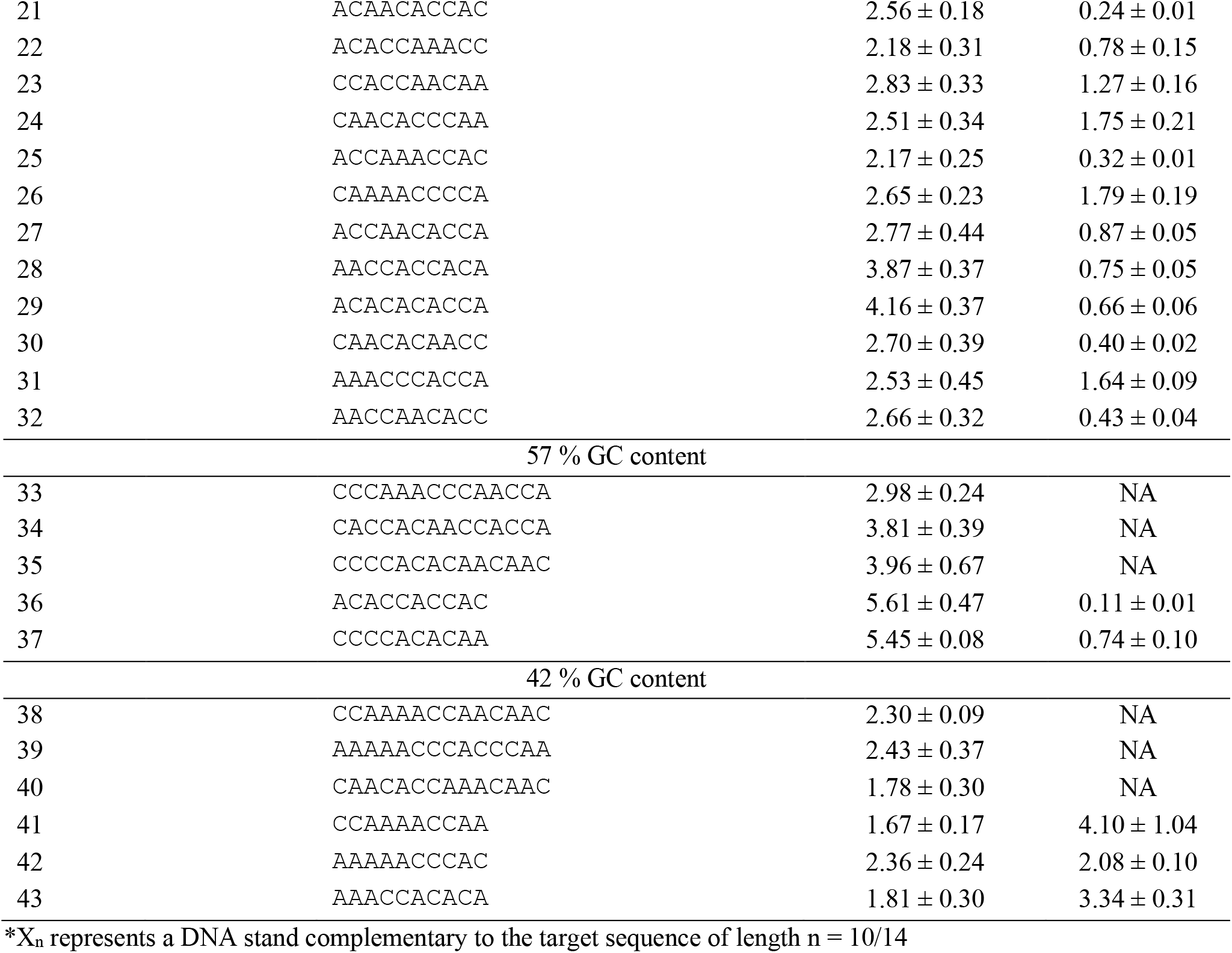
All DNA sequences with associated rate constants measured in this study.

### Surface plasmon resonance experiments

SPR experiments were performed with a Biacore S200 system. A CM5 chip was coated with 4000-5000 RU streptavidin purchased from Sigma-Aldrich. The experimental setup for the surface plasmon resonance measurements was chosen as described before (57), shown in figure 1A. To ensure a Langmuir 1:1 interaction model, an anchor DNA strand was immobilized on the chip surface by biotin-streptavidin coupling at a density of 1.7 × 16^9^ *molecules*/*mm*^2^ so that intermolecular crosslinking of the immobilized DNA strands was minimized. The anchor strand then captured the template strand, which had a free complimentary binding site for the target strand. By referring to the mass and length of the anchor and template strands, the highest signal expected for binding of the target strand to the template was 11 RU (10bp) and 12 RU (14bp), respectively.

**Figure 1:**
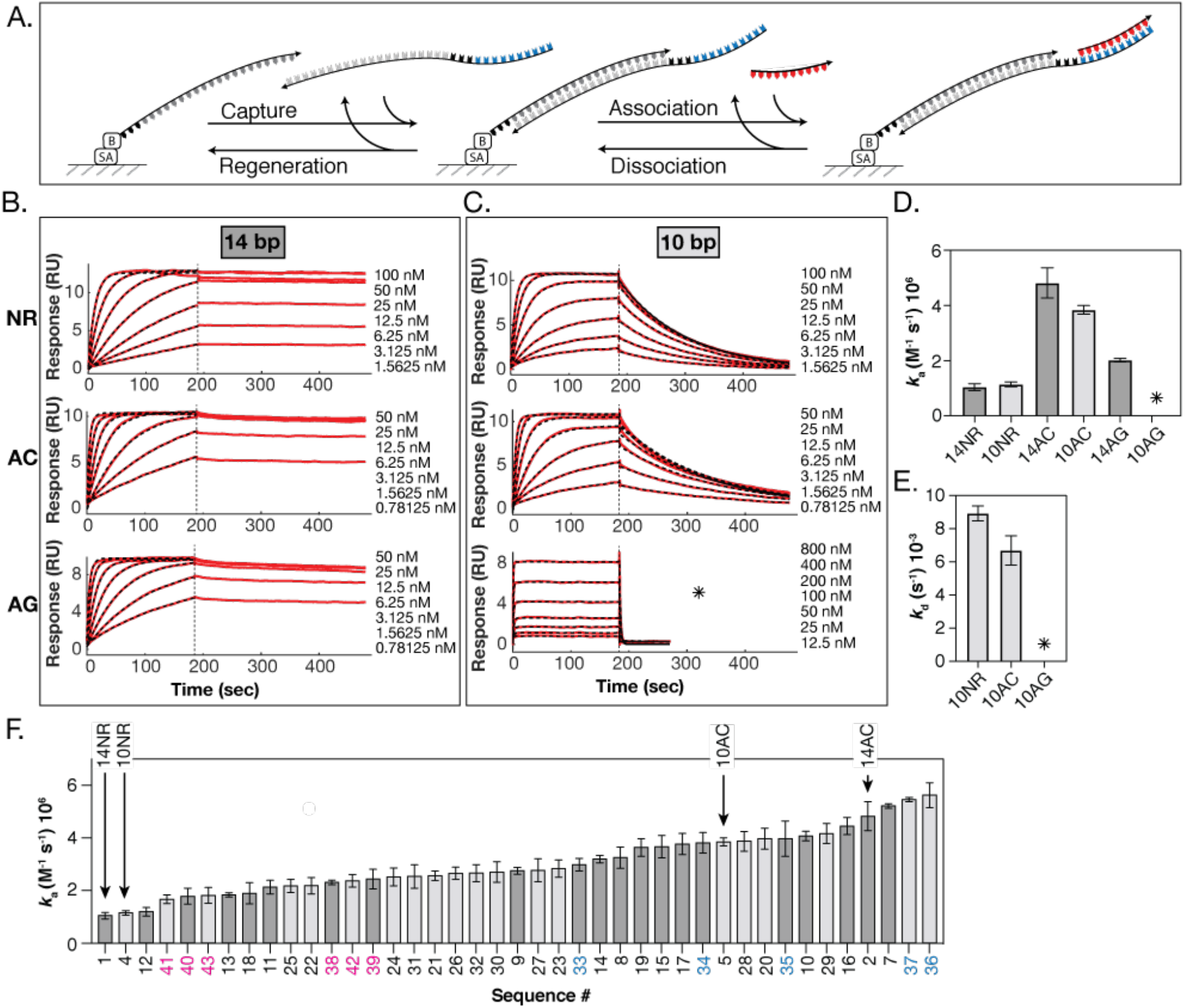
Binding kinetics for non-repetitive and repetitive DNA sequences. (A). Schematic depiction of the surface chemistry used to measure DNA hybridisation kinetics with SPR. First, a 20-base biotinylated DNA strand (anchor, dark grey) binds irreversibly to the streptavidin coated surface of the SPR chip. Second, a longer strand of DNA (template, light grey) that is complementary to the anchor binds (capture). The template strand has an extension consisting of a 3-thymine spacer and a sequence that is complementary to the target strand. Third, association and dissociation kinetics (association and dissociation respectively) of the target strand (red) can then be measured in real time. The chip can be re-used for replicate experiments after a regeneration step that denatures all DNA duplexes leaving only the black anchor strand (regeneration). (B and C) Representative raw SPR sensorgrams (red) with mono-exponential fit (dashed black) to association phase for 14bp sequences (B) and to association and dissociation phase for 10bp sequences, fit locally for each concentration (C). The apparent high association rate of the 10AG sequence was due to the use of high concentration of target strand required to get an appreciable yield, and the fast dissociation, which increases the rate at which the system approaches equilibrium. Replicate data for sequences in (B) and (C) are in figure S1. (D) Association rate constants for 14bp (dark grey) and 10bp (light grey) sequences. (E) Dissociation rate constants for 10bp sequences. * indicates that no kinetic rates could be determined. (F) Association rate constants for all DNA sequences without secondary structure in this study as measured by SPR and indexed according to Table 1. As in (D), light and dark grey correspond to 10 and 14 base sequences respectively. Sequences with 42% and 57% GC content are marked with magenta and cyan labels respectively. All other sequences have 50% GC content. Error bars are standard deviation from at least three independent measurements. Raw SPR sensorgrams fitted with monoexponential equations are in figure S2-3.

The biotinylated anchor strand was immobilized on two flow cells of the sensor chip, leaving two flow cells as blank reference cells. DNA samples were prepared in 10 mM HEPES pH 7.5, 150 mM NaCl, 3 mM EDTA and 0.005 % Tween20 running buffer and SPR experiments were performed in the same buffer at 25 °C and a flow rate of 60 μl/min. The SPR chip could be regenerated for reuse by removing the template strand with a 60 sec injection of 10 mM glycine pH 2.5. Sensorgrams were double-referenced and three repeats of each data set were carried out. The corrected binding curves were fitted with a 1:1 binding model to obtain apparent association constants, *k*_*app*_, and dissociation constants, *k*_*d*_. *k*_*app*_ were plotted as a function of the target concentration and fit to a linear function whose slope corresponded to the association rate constant, *k*_*a*_. *K*_*D*_ from steady state measurements was calculated using the RU_max_ values obtained from binding curve fits, plotted as a function of the target concentration. All data was fit using Prism and MATLAB. Final *k*_a_ and *k*_d_ values are averages of at least three replicates and errors reported are standard deviations.

### Estimation of binding free energies with NuPACK

Binding free energies were determined using NUPACK version 4.0.0.21 with the following parameters: 25 °C, 0.15 M NaCl, material setting to ‘dna2004’ and ensemble parameter to ‘stacking’.

## RESULTS

### Repetitive DNA sequences hybridise more rapidly than non-repetitive sequences

To experimentally test predictions that additional off-register nucleating interactions in repetitive sequences result in faster hybridisation rates, we compared association rates of two 14 base sequences previously analysed in coarse grained MD simulations (50). The first was a non-repetitive sequence (14NR) with 50% GC content that was designed to minimise hairpin formation and off-register interactions with its complementary strand. The second consisted of seven successive AC repeats (14AC), which maintains the same GC content as 14NR but allows for more off-register nucleating interactions. The absence of complementary bases precludes the formation of secondary structure *via* intramolecular base pairing. In addition, we measured hybridisation kinetics of a repeated sequence consisting of seven successive AG repeats (14AG). Unlike the 14AC sequence, the AG repeat sequence has the capacity to form G-quadruplexes including GAGA quartets (58), and GAGAGAGA heptads (59). Thus, the 14AG sequence provided a convenient means to assess the relative impact of secondary structures and the additional off-register nucleation sites in repeated sequences on DNA hybridisation rates. We also performed measurements with variants of the 14-base DNA sequences that were truncated to a length of 10 bases. Details of all DNA sequences used in this study are summarised in Table 1.

DNA hybridisation kinetics were measured with SPR as previously described (57). DNA strands that were complementary to target strands were immobilised to the surface of an avidin-coated SPR chip by hybridisation to biotinylated ‘anchor’ strands (Figure 1A). During association measurements, target strands were flowed over the surface of the chip at fixed concentrations [T] resulting in pseudo-first-order binding kinetics. The response units (RU) from all SPR sensorgrams were therefore well described by the following monoexponential equation:

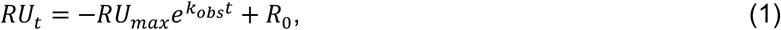

where *RU*_*max*_ is the RU value when all binding sites are occupied and R_0_ is the RU value at the zero time point (Figure 1B and C and S1-3). Association rate constants (*k*_*a*_) could then be calculated from sensorgrams according to:

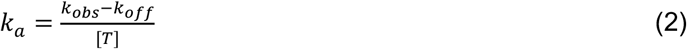

The repetitive 14AC sequence (*k*_*a*_ = 4.8 ± 6.5 × 16^6^*M*^−1^*s*^−1^) hybridised approximately five times faster than the non-repetitive 14NR sequence (*k*_*a*_ = 1.6 ± 6.1 × 16^6^*M*^−1^*s*^−1^) (Figure 1D). This is consistent with MD simulations, suggesting that the additional possible off-register nucleating interactions in repetitive sequences result in faster hybridisation rates (50). As expected, given the propensity for the 14AG sequence to form secondary structures (58, 59), the 14AG sequence hybridised more slowly than the 14AC sequence. Interestingly however, the association rate for the 14AG sequence (*k*_*a*_ = 2.6 ± 6.67 × 16^6^*M*^−1^*s*^−1^) was faster than that of the non-repetitive 14NR sequence, suggesting that additional off-register nucleation states in the AG sequences are sufficient to off-set the reduction of the hybridisation rate associated with the presence of secondary structures. Truncating DNA strands to 10 bases did not appear to have a large effect on association rates. There was no detectable difference in association rates between the 14NR and the truncated 10NR sequence (*k*_*a*_ = 1.2 ± 6.1 × 16^6^*M*^−1^*s*^−1^) and while slower than the 14AC sequence, the 10AC sequence (*k*_*a*_ = 3.8 ± 6.2 × 16^6^*M*^−1^*s*^−1^) still hybridised 4 times faster than the 10NR sequence. Dissociation rates were drastically faster for the AG sequence compared with the AC and NR sequences. The 10AG sequence dissociated too rapidly to be captured within the limits of experimental measurements. Since observed binding curves also depend on dissociation rates (see equation 2), neither association nor dissociation sensorgrams could be fit to monoexponential equations to obtain accurate association or dissociation rates for the 10AG sequence. In contrast, dissociation rates for the 10NR and 10AC sequences were similar and much slower than the 10AG sequence, with a mean dissociation rate of 8.9 ± 6.5 × 16^−2^*s*^−1^ and 6.7 ± 6.9 × 16^−2^*s*^−1^, respectively (Figure 1E). This is consistent with predictions that secondary structures in DNA sequences not only decrease association rates but have a pronounced tendency to increase dissociation rates, possibly originating from the formation of secondary structures during melting (55). It also follows that the relatively slow dissociation rates of the AC and NR sequences reflects the lack of significant secondary structures in these sequences as predicted.

### Association rates of randomly generated AC sequences

DNA sequences consisting only of AC bases (AC sequences) provide a useful means to explore the mechanisms underlying sequence-dependent hybridisation rates in the absence of secondary structure. We therefore measured the hybridisation rates of an additional 38 randomly generated DNA sequences consisting of only adenine and cytosine bases. These DNA strands were either 10 or 14 bases in length with a GC content between 40% and 60%. As above, kinetic traces of all sequences were consistent with pseudo-first-order binding kinetics (Figure S2-3) allowing for reliable determination of binding rates, which are presented in order of increasing rates in figure 1F. The kinetic rate constants for all sequences in this study are summarised in table 1 and supplementary table 1, which also shows, where applicable, consistent equilibrium dissociation constants (*K*_*D*_) as calculated from kinetic rates and steady state measurements, further confirming the reliability of SPR data.

The repetitive 14AC and 10AC sequences were among those with the fastest hybridisation rates ranking 4^th^ and 11^th^ respectively, whereas the non-repetitive 14NR and 10NR had the slowest hybridisation rates (Figure 1F). Furthermore, consistent with previous reports (46), sequences with a higher GC content tended to hybridise more rapidly. Thus, increased hybridisation rates appear to broadly correlate with a greater number and stability of possible nucleating interactions. The dependence of DNA hybridisation rates on strand length may also provide important mechanistic insight. Previous studies are in agreement that dissociation rates significantly decrease with increased DNA strand length. However, data on association rates are mixed with reports of weakly increasing rates (60), decreasing rates (61, 62) or an absence of an effect on the rates (63). Our data also shows no obvious correlation between DNA hybridization rates and sequence length. This suggests either that a length dependent effect on hybridisation rates is insignificant between lengths of 10 and 14 bp or that length related hybridisation properties off-set each other to result in the apparent lack of correlation.

### Simple physically motivated model for capturing the variance in DNA hybridisation rates

To explore the underlying mechanisms dominating DNA hybridisation kinetics in more detail, we constructed a simple, physically motivated model to quantify the correlation between hybridisation rates and the number and stability of nucleation states, including those that are off-register, that result in a fully formed duplex. This simple model is predicated on the idea that in the absence of other factors such as secondary structure, sequence-dependent hybridisation rates are based fundamentally on two factors, the combinatorics of available nucleation sites, and their stability. As illustrated in figure 2, the model assumes that hybridisation proceeds via a nucleation state consisting of a small sub-sequence of *n* contiguous intermolecular base pairing interactions (30, 48-50, 52). *n* thus constitutes a model parameter controlling the effect of combinatorics of the sub-sequences in the strands. From this nucleated state, the strands either dissociate and return to solution or transition into one of a vast number of complicated states associated with various intermediary and meta-stable complexes including partially zippered, off-register structures from where the complex can proceed to a fully hybridised duplex (50).

**Figure 2.**
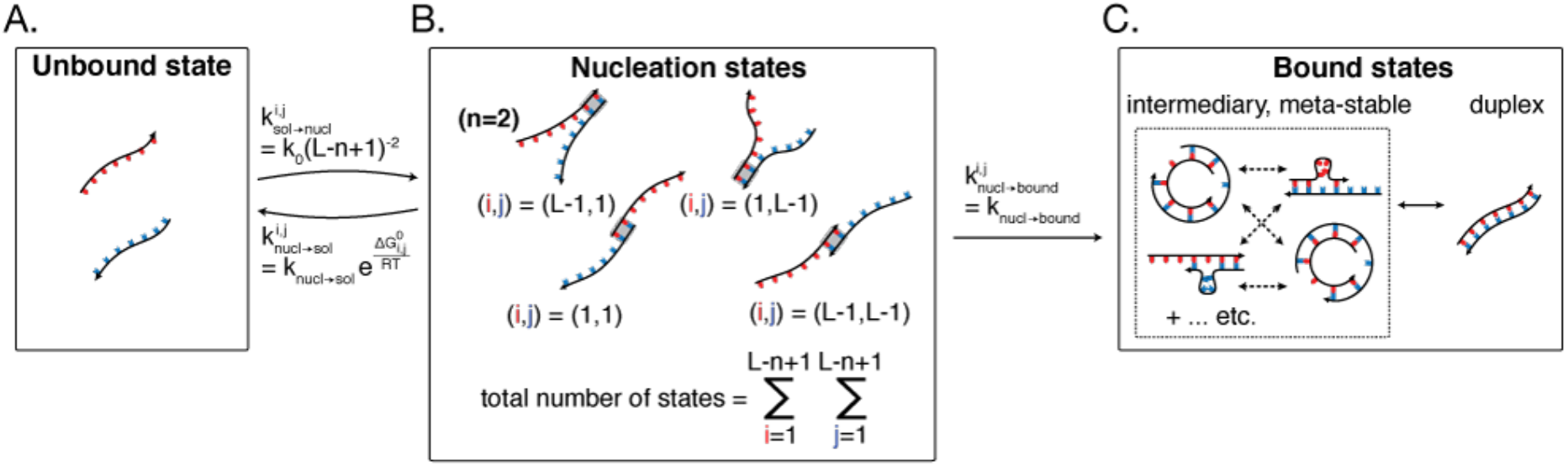
Cartoon depiction of nucleation and hybridisation underpinning the simple model with a nucleation length of two. From the unbound state (A) the system can transition into one of (*L* − *n* + 1)^2^ possible nucleation states, where *L* is the length of the DNA strands and *n* is the length of the nucleating interaction. (B) illustrates examples of the many possible nucleation states of length *n* = 2. From any of these states the system can either return to an unbound state, or it can continue through to full hybridisation via a complicated network of possible intermediary and meta-stable bound states as illustrated in (C). The model assumes negligible transitions from bound states back to nucleated states. Rates are for transitions to and from a single specified location (*i,j*), where *i* and *j* refer to the index of the first base involved in the nucleating interaction from the 5’ end, on each strand respectively. For each possible nucleation location there exists a constant rate of nucleation equal to R _κ_ (*L* − *n* + 1)^−2^ such that the overall rate of nucleation is independent of length. Then there is a sequence dependent rate from the nucleated state back to solution that depends on the stability of the nucleated binding complex. Finally, there is a constant rate of transitioning from the nucleated state into a bound state. From these rate definitions, the effective rate of hybridisation due to nucleation location (*i,j*) is taken as the inverse of the mean first passage time from the solution state to the bound state (equation (6), Supplementary Note 1).

If transitions from an unbound state into a nucleated state are rate-limiting, and progression to eventual full hybridisation from a meta-stable structure is very likely (50), then a faithful description of the kinetics between the unbound, nucleation and meta-stable states will provide an approximate measure of the total observed rate of hybridisation. We can thus construct a framework for modelling the effective hybridisation rate constant using the simple form:

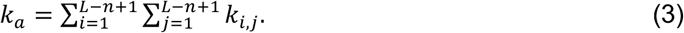

Here, *L* is the length of the strand expressed as an integer number of bases, whilst *i* and *j* are indices corresponding to the position of the first of *n* contiguous bases which make up the nucleation state, in the 5’ → 3’ direction, for the strand and its complement, respectively (Figure 2B – top). The double sum therefore includes (*L* − *n* + 1)^2^ contributions from all such nucleation states, regardless of whether they are on-register, or whether they are formed from complementary bases (Figure 2B - bottom). *k*_*i,j*_ then quantifies the specific contribution arising from the nucleation state at positions *i* and *j* on the strand and its complement, respectively. Crucially, unlike previous models (45), this allows the contribution of any particular nucleation state to vanish in the case of mis-matched bases, thus naturally capturing the combinatorics of nucleating interactions, and for the contribution of nucleating interactions to vary with the stability of the nucleated state when they do match.

To account for the relative stability of each *i*− *j* nucleation site we can approximate the associated contributing rate constant *k*_*i,j*_ as arising from an idealised sub-system consisting of free or dissociated strands in solution, a single *i,j* nucleation state, and a ‘bound’ state representing all configurations where the strands are in one of many more complicated subsequent complexes including fully hybridised duplexes or pseudoknots etc. (Figure 2C). Thus, for any given individual *i,j* nucleation site, this three state sub-system is then fully described with the specification of the rate constants associated with transitions between these states (Figure 2). Various assumptions can then be implemented to define the rates at each step.

First, we specify a transition rate from a nucleated state to dissociated strands in solution, which we capture as a stability term measured through the free energy of binding of the nucleation state, and thus introducing explicit sequence dependence into the model,

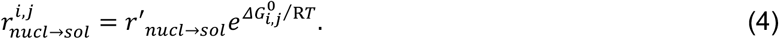

Here 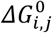 is the free energy of binding of the nucleation state associated with binding locations *i* and *j* measured in J/mol, *r*′_*nucl*→*sol*_ is a rate constant that is independent of the binding sequence, R is the gas constant and T is the temperature in Kelvin. Second, we ignore any entropic effects of unbound DNA bases surrounding the nucleation site and assume that all specific nucleation sites are equally accessible from dissociated strands. The rate for forming any given *i, j* nucleating interaction is assumed to be constant across all nucleation sites, and can be given by 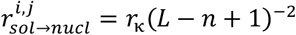 where R _κ_ is an overall scaling factor representing the rate of a nucleation event occurring in any pair of locations on the strands independently of their length, comprising a rate constant κ and any concentration dependence (e.g. R _κ_ = κ [*T*] in the pseudo-first-order conditions above), and the (*L* − *n* + 1) ^−2^ term imposes the observed lack of scaling of hybridisation rates with strand length on the model. Third, we assume that the rate of transitions from the bound state to the nucleated state is slow relative to the time scales of hybridisation and hence these transitions are ignored in the model. Finally, as a first approximation, we assume that the microscopic rate of transition from any nucleated site into an intermediary or metastable state is constant across all nucleation sites and sequences 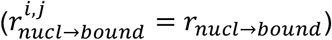.

The effective rate for hybridisation via any given *i,j* nucleating interaction can be arrived at by computing the inverse of the mean first passage time taken to transition from state 1 to state 3. From the rates defined above this rate is given by (Supplementary Note 1):

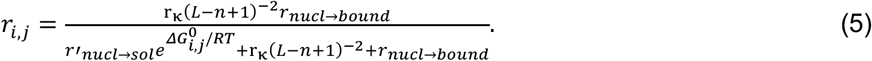

While nucleation is the rate limiting step, the rate of transitions away from the nucleated state are much faster than the rate of transitions into the nucleated state (*r*_*nucl* → *bound*,_ *r*′_*nucl* →*sol*_ ≫ r _κ_ (*L* − *n* + 1*x*^−2^). As such we can simplify this expression, to leading order in R _κ_ (*L* − *n* + 1)^−2^/ *k*_*nucl*→B*ound*_, and then convert to the relevant rate constant to find

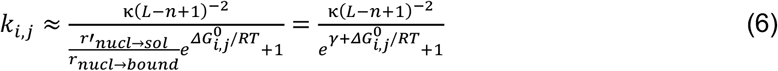

where *γ* = ln (*r* ′_*nucl*→*sol* /_ *r*_*nucl*→B*ound*_. The rate constant for hybridisation via any single (*i,j*) nucleation state can thus be directly interpreted as a uniform and limiting rate, κ(*L* − *n* + 1)^−2^, into the nucleation state from solution multiplied by a probability of continuing through to full hybridisation from the nucleation state,

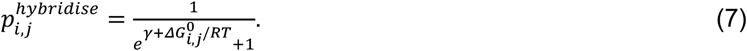

Consequently, the value of *γ* coincides with the value of 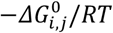 for which the probability of continuing on to hybridisation is 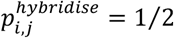. Substituting equation (6) into equation (3) we arrive at

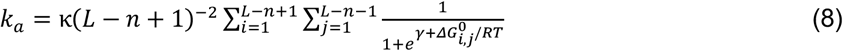

fully specifying our model up to estimation of the nucleation free energies of the nucleation state. When a nucleation state (*i,j*) constitutes a mismatch the model considers the nucleation free energy to be infinity such that the probability of hybridisation is zero. For complementary nucleation states, nucleation free energies, 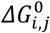 were obtained using the NUPACK 4.0.0.21 implementation of the nearest neighbour model (see materials and methods).

Given a fixed *n*, the terms κ and *γ* then constitute the free parameters of the model, which can be fit to data. However, of the two, only *γ* controls the sequence dependence, with κ simply acting as a scaling factor. In physical terms *γ* controls how sharply the increases in the stability of the nucleation states increases the likelihood of continuing through to full hybridisation. Crucially, the fact that the sequence dependent stability of nucleating interactions is controlled by a single free parameter dramatically restricts model complexity such that over-fitting can be avoided as much as possible.

### Fits to experimental data

We first determine how well nucleation site combinatorics alone correlate with relative hybridisation rates, without accounting for stability. This is achieved simply with the model described above (Equation 8) by replacing the probability of hybridisation with a value of one if the nucleation site is formed from complementary base pairs, or zero with mismatched base pairs. All other aspects of the model are unchanged. The resulting model rates can then be scaled to fit with experimental data by varying the scale factor, κ, using a Nelder-Mead optimisation algorithm taking the sum of the squared residuals as the objective function to be minimised. Fits were performed with nucleation lengths of *n* = 1,2,3 *and* 4 and predicted rates plotted against measured rates (Figure 3A) from which correlation coefficients were calculated. Given the complexity of DNA hybridisation, and the potential that many rate determining factors were not accounted for in our simple model, it was important to ensure that reported correlations were a true reflection of model accuracy. We therefore performed careful statistical analysis to estimate standard deviations and confidence intervals to quantify the certainty in correlation coefficients. In addition, we performed permutation tests to determine p-values based on the probability that similar correlations could be obtained from null-distributions of randomly reshuffled datasets. Where correlations between model and experiment were high (*ρ* > 0.5) p-values were low (*p* < 6.661), thus providing confidence that the observed trends were not spurious. Further, null distributed correlations were very close to 0 (Supplementary Table 2) providing confidence that the observed correlations were not due to overfitting. A detailed description of error analyses performed in this study is in supplementary note 2. Standard deviations, confidence intervals and p-values for all reported correlation coefficients are in supplementary table 2, and model parameters from all fits in this study are in supplementary table 3. For completeness, we also report correlation coefficients calculated from point values in supplementary table 4, which do not account for experimental uncertainty.

**Figure 3.**
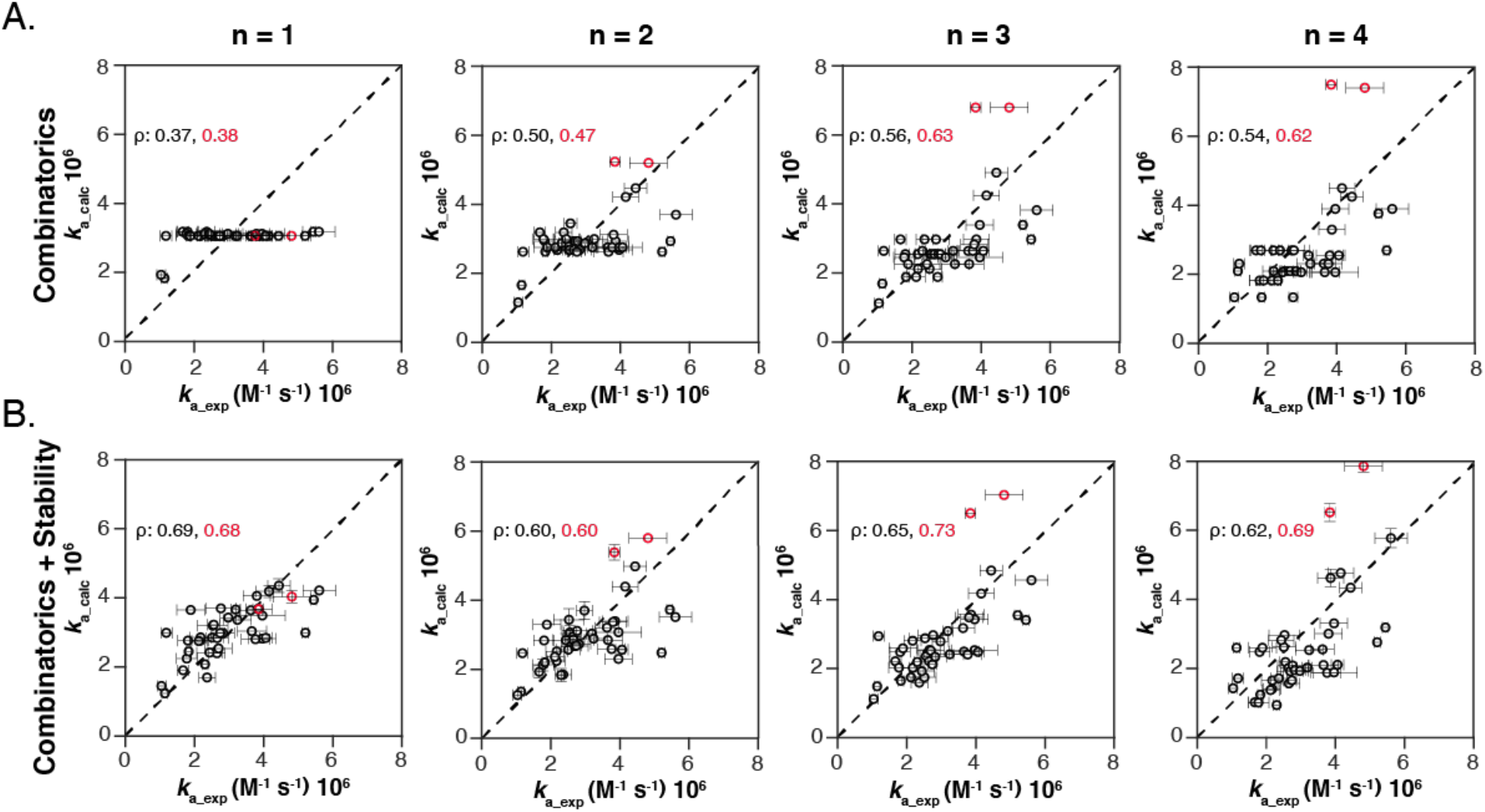
Predicted vs measured hybridisation rates. Predicted rates from combinatorics alone in (A) and from the full model in (B) with *n* = 1 to *n* = 4 from left to right. Errors are standard deviations from at least three independent measurements. Red data points depict rates for repetitive 10AC and 14AC sequences. Each plot is labelled with correlation coefficients for the entire dataset (black) and for data omitting the repetitive 10AC and 14AC sequences.

With a nucleation length of *n* = 1, combinatorics alone provides no distinguishing power between different AC sequences with the same GC content, since these sequences necessarily have the same number of possible complementary one base interactions. Consequently, predicted rates using combinatorics alone were essentially flat for AC sequences when *n* = 1 (Figure 3A). Additionally, the NR sequences, which also consist of G and T bases, can make far fewer complementary single base pair interactions and thus have lower predicted hybridisation rates, consistent with experiment. Indeed, any weakly existing correlation when *n* = 1 can be attributed to the slower hybridisation rates for the NR sequences. With increasing nucleation length, there is a larger variation in the number of complementary nucleating interactions. This variation yields a clearly positive correlation between predicted and measured hybridisation rates (Figure 3A) that increases with nucleation length reaching a maximum when *n* = 3, which has a correlation coefficient of *ρ* = 6.56 ± 6.64.

To incorporate base-pair stability in the model, we next fit the full model in equation (8) to experimental data to determine whether improved fits could be obtained over predictions based on nucleation site combinatorics only. Accounting for stability introduces the single free parameter, *γ*, which along with the scaling parameter κ, was varied, again taking sum of the squared residuals as the objective function to be minimised for each nucleation length (*n* = 1−2−3 *or* 4*x*. Relative to combinatorics alone, the full model resulted in an improved correlation between model and experimentally measured hybridisation rates across all nucleation lengths (Figure 3B). We note in particular a high correlation with a nucleation length of *n* = 1 (*ρ* = 6.68 ± 6.63), where there was a lack of correlation from combinatorics alone and which therefore can be attributed almost exclusively to the inclusion of nucleation site stability.

For higher binding lengths, *n* > 1, stability reliably improves the achieved correlation, but none attain the correlation captured by the n=1 case. We observe however that the model consistently over-estimates the related, and most repetitive sequences, 10AC and 14AC, possibly indicative of a lurking feature limiting the increase of hybridisation rates due to increasing combinatorics not captured by the model. If we omit these two related sequences the maximal correlation between predicted and measured hybridisation rates is improved to *ρ* = 6.73 ± 6.63, occurring at *n* = 3 (Figure 3B and S4 and Supplementary Table 2).

## DISCUSSION/CONCLUSION

This study explores the complex processes underpinning DNA hybridisation and sequence-dependent binding kinetics. While previous studies identified secondary structure as a key contributing factor to hybridisation rates (45, 56), we focus on other, equally relevant but poorly defined physical factors to gain a more complete understanding of DNA hybridisation. In particular, by combining careful experimental design and measurements with a physically justified theoretical model, significant progress is made in cementing several principles, proposed to be fundamental to DNA hybridisation mechanisms. These principles are: that the rate of forming nucleating interactions limits the rate of DNA hybridisation (30, 48, 49); that the combination and stability of all possible nucleating interactions is therefore a rate determining factor (45, 50); and that rate-limiting nucleating interactions can be off-register from a fully formed duplex (50). The study experimentally verifies previous predictions that repetitive sequences, which have a greater number of off-register nucleating interactions, hybridise more rapidly than non-repetitive sequences (50). Additional measurements were then performed to capture the variance in hybridisation rates between 41 different strands that have little or no secondary structure (Figure 2). Finally, a simple physically motivated model has been developed that captures a large part of this variance by accounting only for the combination of possible nucleating interactions, including those that are off-register, and their relative stability.

A guiding principle in the construction of our model is to use as few free parameters as possible, to avoid over-fitting and enable the inference of broad physical mechanisms in the context of limited and/or noisy data. However, such a principle is always in tension with a faithful representation of the complicated physical processes that underpin DNA hybridisation. We sought to achieve a useful balance by focussing on two physically plausible rate determiners: the combinatorics and thermodynamic stability of nucleation states. Our model in turn captures these factors with a minimal number of parameters, the nucleation length *n* and the stability term *γ*. We have purposely avoided introducing additional confounding factors as much as possible with the exclusive focus on DNA sequences that exhibit very little to no secondary structure. A consequence of this approach is that we do not expect, nor is it our intention, that we will perfectly capture all the observed variance in experimental hybridisation rates. Instead, our goal was to capture as much variance as possible by modelling physically plausible mechanisms with as few parameters as possible. Consequently, the strong correlation achieved between model and experimental data (*ρ* = 6.73 ± 6.63) provides valuable insight into the physical mechanisms of DNA hybridisation and sequence dependent hybridisation rates.

Our approach is in contrast to several recent studies. For instance, Hata *et al*. achieved a very good correlation with experimental data using a model with a physically motivated component based on secondary structure. However, the high correlation is contingent on an additional sequence dependent component involving 32 free parameters which are difficult to interpret physically (45). Other studies lean more heavily on data driven methodologies, again obtaining reasonable correlations, but requiring the use of a large number of data features. These studies thus provide little insight into physical mechanisms apart from secondary structure, that may be underpinning the process.

The repetitive 10AC and 14AC sequences had the greatest combination of nucleating interactions, particularly at longer nucleation lengths, which resulted in model predictions that were substantially higher than experiment, but only when *n* ≥ 3 (Figure 3). Indeed, the results of the model are highly contingent on the choice of the binding nucleation site length, *n*. Moreover, there was no single choice of nucleation length (*n* = 1,2,3 *or* 4) that yielded correlations that were drastically better than the others. Here we must emphasise, the restriction to a single nucleation length is highly idealised whereas in reality, nucleation is a progressive and complicated process. Microscopically, a practically innumerable number of nucleation and hybridisation pathways will exist from free strands to full hybridisation, and these pathways will be distinct for different initial interactions between the strands. Along different parts of these pathways further progression will be practically guaranteed, whilst at others it may be highly unlikely. As such, the success of any given binding nucleation length does not imply importance to the exclusion of other characteristic lengths, but simply reflects that it is possible to capture the variance in the data by examining an ostensibly critical part of the nucleation process. In this respect one can view the choice of the nucleation state being *n* bases as constituting the implicit assumptions that any nucleation state with fewer than *n* bases will disassociate if they cannot lead to a contiguous nucleation state of *n* bases, and that binding states of more than *n* contiguous bases are very likely to lead to full hybridisation. In reality however, it is unlikely that there exists a single threshold nucleating length that fulfils the criteria above. Indeed, plotting the distribution of binding free energies at different *n* shows considerable overlap in the distribution of binding free energies at different nucleation lengths (Figure S5). These distributions suggest, according to NUPACK predictions, that a larger *n* does not necessarily translate to a more stable nucleating interaction. Thus, the true threshold length of a nucleating interaction according to the definition above, is highly sequence dependent and will differ not only between DNA strands with different sequences but also within any given DNA strand. One could extend this model to account for variable nucleation lengths, but this would unavoidably lead to a proliferation of associated free parameters, drastically increasing the chance of over-fitting.

The probability that any matched nucleating interaction will proceed towards a fully formed duplex, can be calculated from the stability of nucleating interactions (as determined by NUPACK) combined with the *γ* value obtained from fits to experimental data according to equation (7). These probabilities for all such interactions in this study are reported in model outputs, which are available in a GitHub repository, with a mean probability of 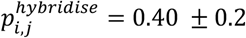, which is similar to previous observations from MD simulations (50). However, model probabilities rely on the stipulation that there is a constant rate of progression from an initial nucleation state to more strongly bound metastable or fully hybridised states, which is also a simplification of the real underlying physical hybridisation process. Attempting to account for such variation would again inevitably introduce large numbers of additional free parameters, risking over-fitting which, even if appropriately implemented, may in turn serve only to obscure more primary physical principles behind the variation of hybridisation rates.

The hybridisation rates of 10 base and 14 base DNA strands were similar with no obvious correlation between sequence length and hybridisation rates (Figure 2I), which informed the decision to normalise the model prediction by the number of possible nucleation states, (*L* − *n* + 1)^−2^, thus removing length dependence from our model. For completeness however, we also performed optimisations with the negative square replaced by a free exponent, *αi*. Fitted values of *αi* that are less than or greater than -2 would be suggestive of a positive or negative dependence of hybridisation rates with strand length. However, the use of such an additional parameter conferred very little increase in model performance, with optimised values of *αi* very close to -2 (Supplementary Table 2), as emerges from the initial model choice and confirming that the dependence on length in our data is extremely weak. While a weak dependence of hybridisation rates on length cannot be expected to hold true for all lengths of DNA strands, the lack of length dependence in our data could be the result of many possible physical processes. A natural interpretation in terms of the presented model is that the number of nucleation attempts per unit time were constant across sequence lengths, perhaps due to similar effective diffusion coefficients and molecular cross-sections over the range of lengths utilised. In turn this property may be contingent on the designed lack of secondary structure in our data set.

Despite these strong assumptions, our model has favourable properties, which strengthen its claims for a faithful capturing of basal physical processes. First, the model is constructed from minimal free parameters, and second, the variation possible in the model is strongly constrained by physical plausibility arguments. Consequently, the capacity for fitting arbitrary patterns in data is severely constrained. Explicitly, all other factors being equal, the model always assigns greater hybridisation rates to sequences that have a larger number of repetitive sub-sequences and when the stability of those binding states is stronger, with the sole variation controlling the size of such an effect. If some other physical property were more dominant, which conflicted with the property that more stable nucleation sites hybridise faster for example, the model would be unable to capture it. Thus, while our model cannot be taken as a precise account of the hybridisation process, it enables us to conclude the correlations it achieves with data lends strong evidence to the claim that both binding site combinatorics and the stability of those sites are strongly implicated as dominant mechanisms underlying the sequence-dependent hybridisation rates of DNA strands *in vitro*. These findings will be useful for the design of applications in DNA nanotechnology such as DNA PAINT where control over hybridisation kinetics is imperative for achieving adequate signal to noise within practical acquisition times (28, 64, 65) (66, 67). Future work could incorporate our findings with approaches such as by Hata *et al*. (45), whose algorithm explicitly accounts for the consequences of secondary structure on nucleation propensities.

## DATA AVAILABILITY

Code for fitting the model to experimental data, all model outputs, and binding energies for all nucleating interactions for all DNA sequences in this study are available in the GitHub repository (https://github.com/llee0905/DNA-bind)

## SUPPLEMENTARY DATA

Supplementary Data are available at NAR online. These include Supplementary Figures and Supplementary Tables in Supplementary Data File 1 and Supplemental Notes in Supplementary Data File 2.

