## Supplementary Information for "Mechanisms underlying sequence-dependent DNA hybridisation rates in the absence of secondary structure"

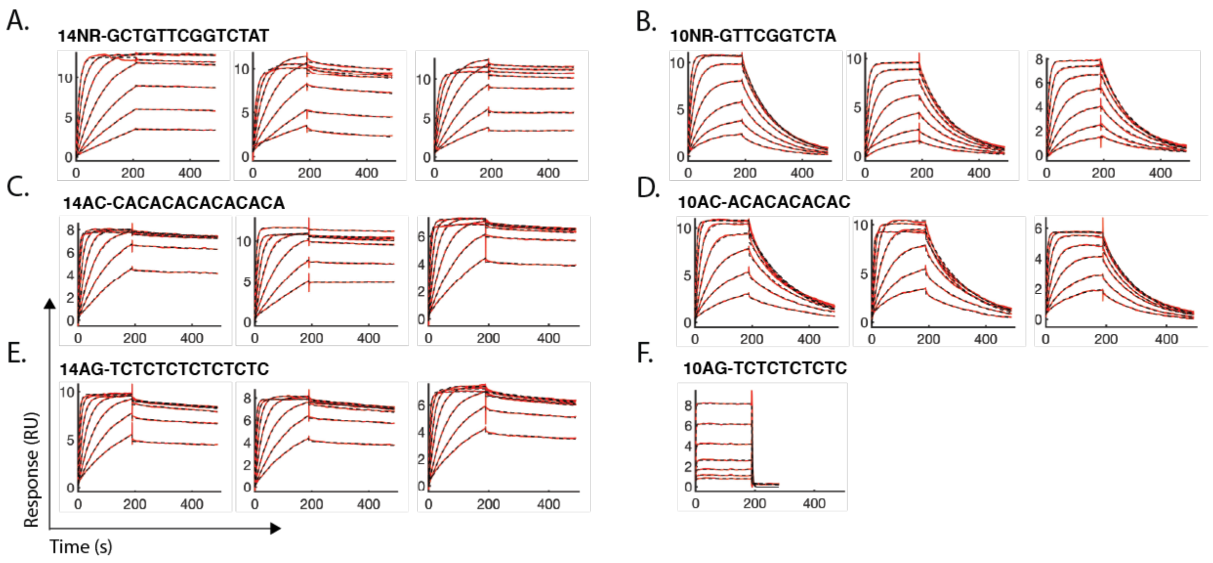


**Figure S1. Replicate SPR sensorgrams for repetitive AC and non-repetitive sequences.** Raw sensorgrams (red) with mono-exponential fits for association and dissociation (dashed black) are shown for sequences associated with Figure 1 in main text. Curves are serial dilutions from 100 nM to 1.5625 nM (A,B) 50 nM to 0.78125 nM (C-E) and 800 nM to 12.5 nM (F).

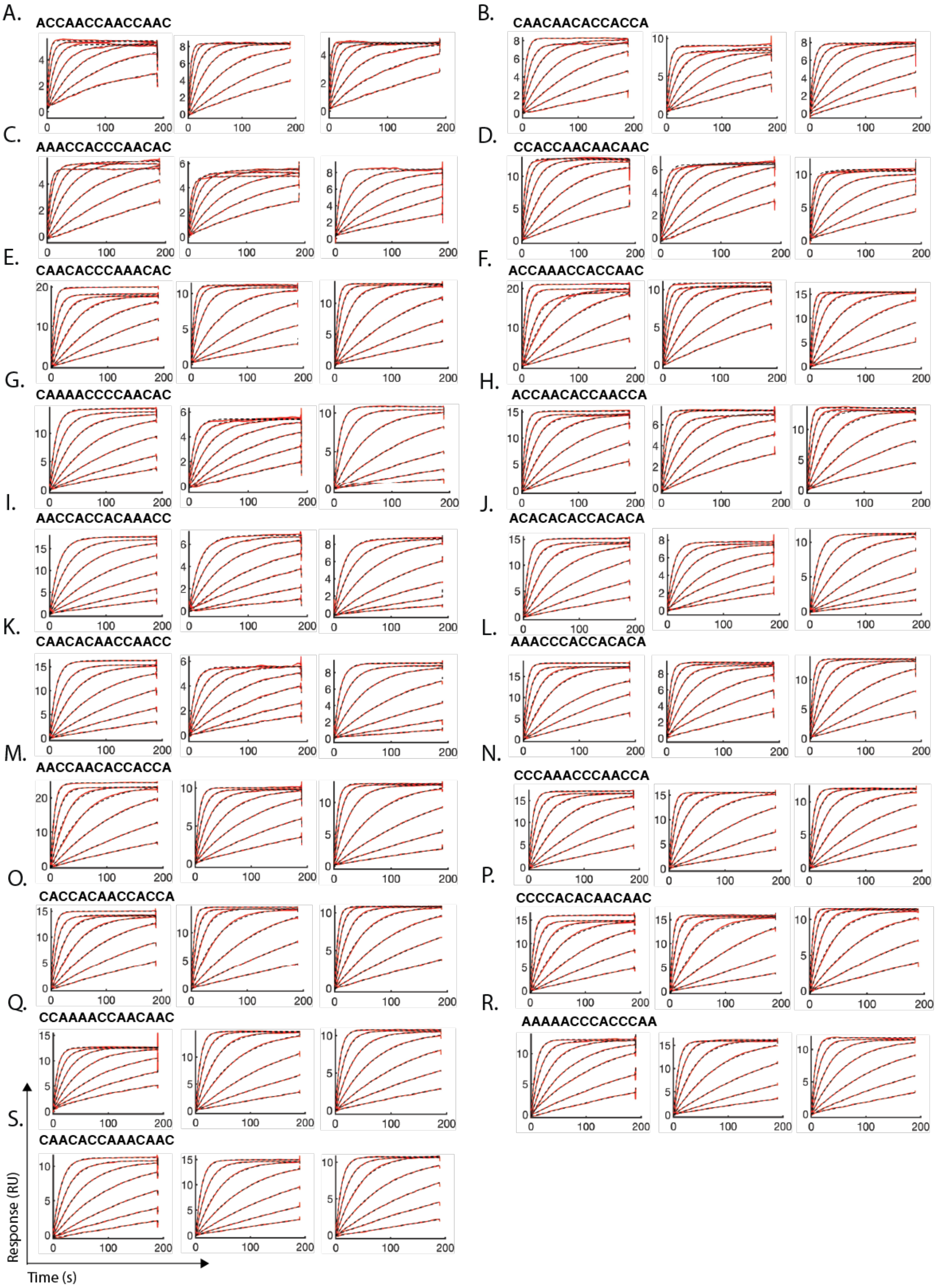


**Figure S2. Replicate SPR sensorgrams for all non-repetitive 14 base AC sequences.** Raw sensorgrams are shown in red with mono-exponential fits overlayed in dashed black). Concentrations are serial dilutions from 50 nM to 0.78125 nM.

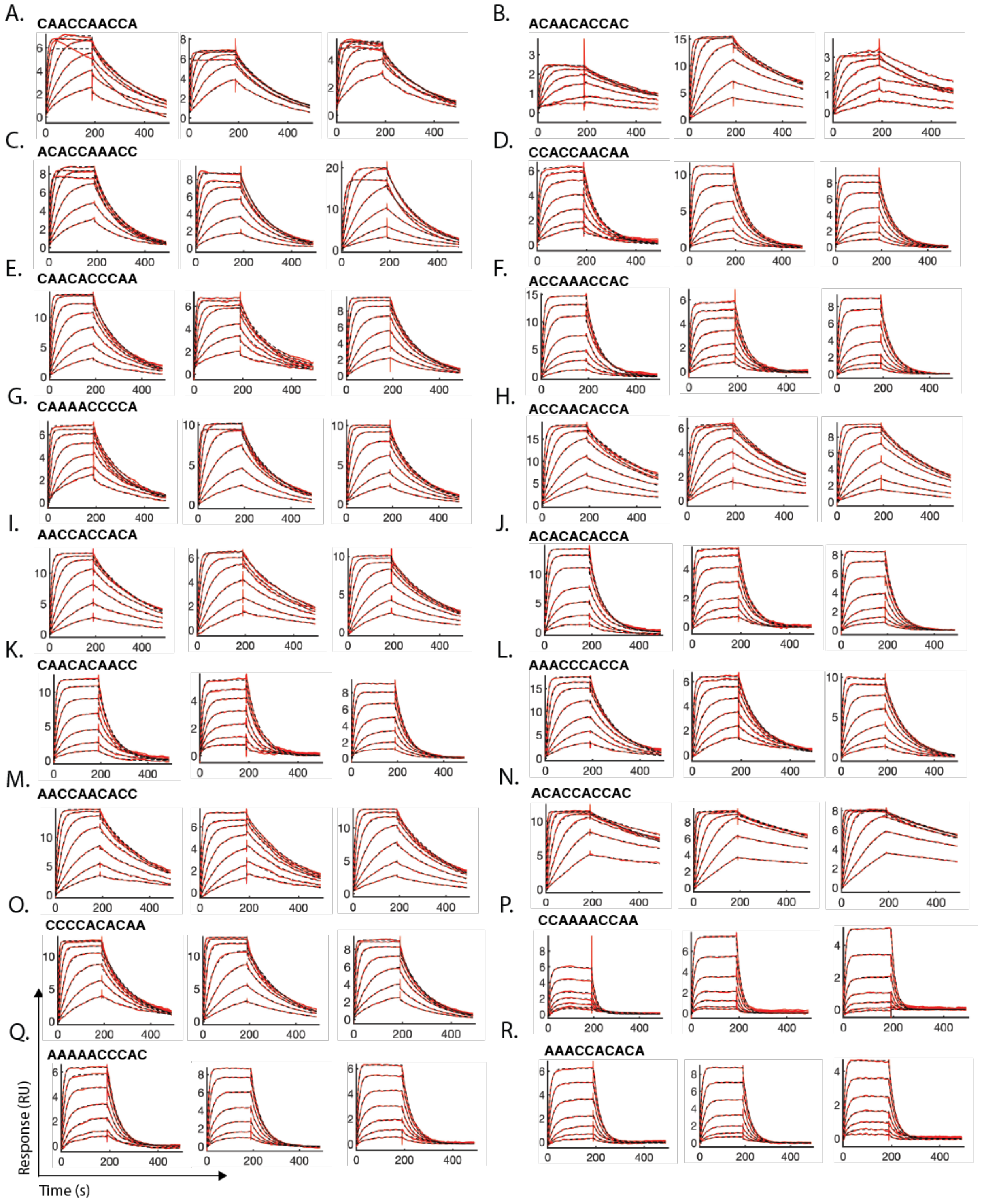


**Figure S3. Replicate SPR sensorgrams for all non-repetitive 10 base AC sequences.** Raw sensorgrams are shown in red with mono-exponential fits for association and dissociation overlayed in dashed black). Concentrations are serial dilutions from 50 nM to 0.78125 nM.

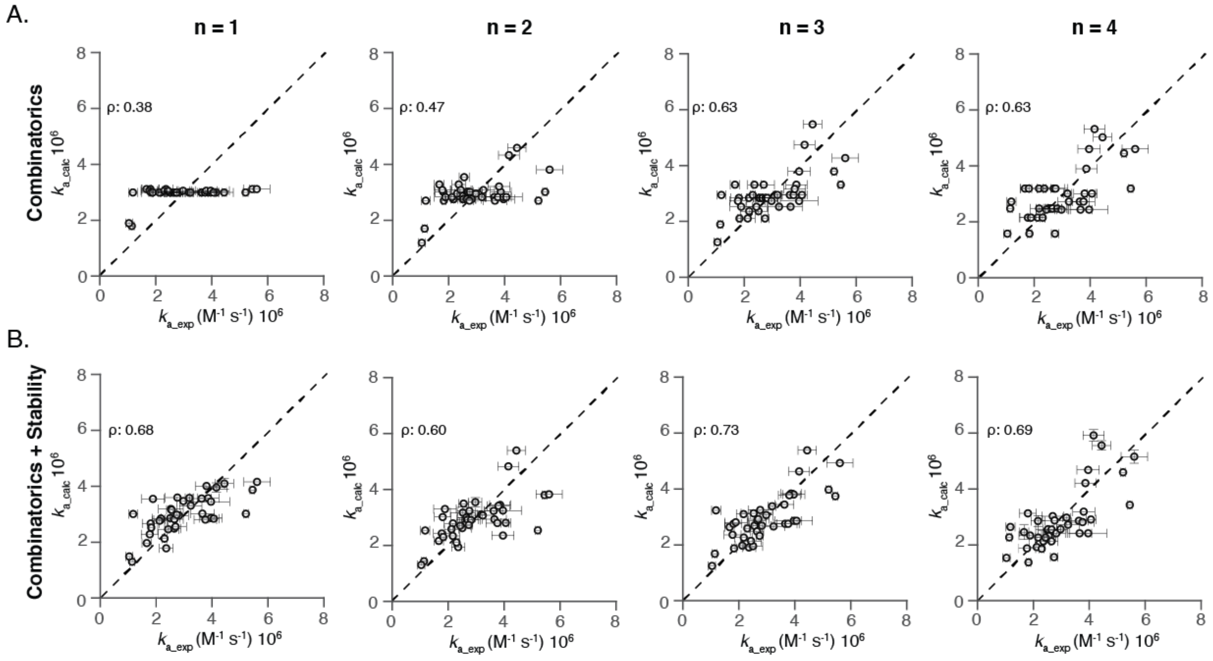


**Figure S4. Model vs experimentally measured hybridisation rates without repetitive 10AC and 14AC sequences.** (A) shows fits from combinatorics alone and (B) from the full model, which includes stability. Errors are standard deviations from at least three independent measurements.

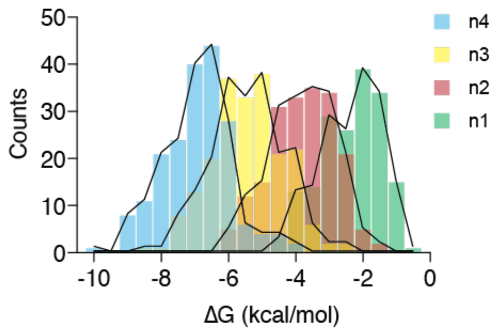


**Figure S5. Distribution of binding energies at different nucleation lengths for all unique nucleation states in this study.**

Supplementary Table 1: Comparison of equilibrium dissociation constants determined from kinetic rate constants and from steady state measurements for all 10 base DNA sequences in this study.

| Number | Name | Sequence 5’ > 3’ | $K_D$ (nM) kinetics | $K_D$ (nM) steady state |
|---|---|---|---|---|
| 4 | 10NR | GTTCGGTCTA | 7.7 ± 0.2 | 6.7 ± 0.1 |
| 5 | 10AC | ACACACACAC | 1.6 ± 0.1 | 1.3 ± 0.2 |
| 6 | 10AG | TCTCTCTCTC | NA | 212 ± 15 |
| | | 50 % GC content | | |
| 20 | | CAACCAACCA | 1.4 ± 0.1 | 1.4 ± 0.2 |
| 21 | | ACAACACCAC | 0.9 ± 0.02 | 0.6 ± 0.06 |
| 22 | | ACACCAAACC | 3.7 ± 0.3 | 3.9 ± 0.4 |
| 23 | | CCACCAACAA | 5.1 ± 0.9 | 6.3 ± 1.8 |
| 24 | | CAACACCCAA | 7.0 ± 0.4 | 7.8 ± 3.2 |
| 25 | | ACCAAACCAC | 1.5 ± 0.2 | 1.4 ± 0.4 |
| 26 | | CAAAACCCCA | 6.8 ± 0.6 | 7.1 ± 2.3 |
| 27 | | ACCAACACCA | 3.2 ± 0.5 | 3.1 ± 1.1 |
| 28 | | AACCACCACA | 2.0 ± 0.2 | 1.9 ± 0.3 |
| 29 | | ACACACACCA | 1.7 ± 0.1 | 1.5 ± 0.1 |
| 30 | | CAACACAACC | 1.5 ± 0.2 | 1.4 ± 0.05 |
| 31 | | AAACCCACCA | 6.6 ± 0.8 | 5.8 ± 0.7 |
| 32 | | AACCAACACC | 1.6 ± 0.1 | 1.4 ± 0.5 |
| | | 57 % GC content | | |
| 36 | | ACACCACCAC | 0.2 ± 0.03 | NA |
| 37 | | CCCCACACAA | 1.3 ± 0.2 | 1.4 ± 0.4 |
| | | 42 % GC content | | |
| 41 | | CCAAAACCAA | 24.9 ± 6.3 | 28.1 ± 5.1 |
| 42 | | AAAAACCCAC | 8.9 ± 0.6 | 8.4 ± 1.8 |
| 43 | | AAACCACACA | 18.8 ± 3.8 | 16.0 ± 4.8 |

**Supplementary Table 2:** Statistics of the correlation coefficient, $\rho$, between experimental data and model predictions, for the null distribution sampled by random permutations, and resultant statistics of $p$-value estimates.

| Model | n | Outliers excluded† | Data | | Null | | $p$-value | |
|---|---|---|---|---|---|---|---|---|
| | | | mean (std-dev) | 95% CI | mean (std-dev) | 95% CI | mean (std-dev) | 95% CI |
| Combinatorics | 1 | N | 0.368 (0.0189) | [0.332, 0.406] | $-4.71\times10^{-5}$ (0.158) | [-0.329, 0.279] | 0.00223 (0.00169) | [0.000250, 0.00654] |
| Comb. + length | 1 | N | 0.371 (0.0187) | [0.335, 0.408] | 0.0777 (0.161) | [-0.262, 0.362] | 0.0218 (0.00819) | [0.00920, 0.0411] |
| Stability | 1 | N | 0.688 (0.0254) | [0.636, 0.736] | 0.0508 (0.165) | [-0.295, 0.345] | $1.02\times10^{-5}$ ($5.53\times10^{-6}$) | [$5.13\times10^{-6}$, $2.50\times10^{-5}$] |
| Stab. + length | 1 | N | 0.690 (0.0253) | [0.639, 0.738] | 0.118 (0.163) | [-0.227, 0.406] | $1.39\times10^{-5}$ ($1.18\times10^{-5}$) | [$5.19\times10^{-6}$, $4.30\times10^{-5}$] |
| Combinatorics | 2 | N | 0.495 (0.0364) | [0.421, 0.563] | 0.000317 (0.158) | [-0.300, 0.315] | 0.00107 (0.00115) | [$7.79\times10^{-5}$, 0.00406] |
| Comb. + length | 2 | N | 0.504 (0.0384) | [0.426, 0.576] | 0.0399 (0.162) | [-0.271, 0.359] | 0.00205 (0.00215) | [0.000184, 0.00781] |
| Stability | 2 | N | 0.600 (0.0363) | [0.525, 0.667] | 0.0554 (0.160) | [-0.256, 0.365] | 0.000192 (0.000305) | [$8.58\times10^{-6}$, 0.000999] |
| Stab. + length | 2 | N | 0.602 (0.0367) | [0.527, 0.670] | 0.0886 (0.162) | [-0.230, 0.400] | 0.000447 (0.000623) | [$2.78\times10^{-5}$, 0.00207] |
| Combinatorics | 3 | N | 0.559 (0.0389) | [0.478, 0.630] | 0.000282 (0.158) | [-0.292, 0.320] | 0.000257 (0.000405) | [$1.61\times10^{-5}$, 0.00132] |
| Comb. + length | 3 | N | 0.569 (0.0414) | [0.484, 0.646] | 0.0278 (0.162) | [-0.273, 0.352] | 0.000391 (0.000666) | [$1.31\times10^{-5}$, 0.00215] |
| Stability | 3 | N | 0.655 (0.0366) | [0.578, 0.721] | 0.0158 (0.159) | [-0.279, 0.335] | $2.85\times10^{-5}$ ($3.87\times10^{-5}$) | [$5.96\times10^{-5}$, 0.000105] |
| Stab. + length | 3 | N | 0.658 (0.0371) | [0.580, 0.725] | 0.0428 (0.162) | [-0.262, 0.366] | $4.07\times10^{-5}$ ($9.42\times10^{-5}$) | [$5.39\times10^{-6}$, 0.000263] |
| Combinatorics | 4 | N | 0.543 (0.0397) | [0.461, 0.616] | 0.000473 (0.158) | [-0.292, 0.320] | 0.000412 (0.000595) | [$2.19^{-5}$, 0.00202] |
| Comb. + length | 4 | N | 0.563 (0.0437) | [0.474, 0.645] | 0.0242 (0.161) | [-0.275, 0.349] | 0.000441 (0.000764) | [$8.55\times10^{-6}$, 0.00249] |
| Stability | 4 | N | 0.617 (0.0397) | [0.534, 0.689] | 0.0454 (0.161) | [-0.263, 0.362] | 0.000134 (0.000268) | [$7.97\times10^{-6}$, 0.000776] |
| Stab. + length | 4 | N | 0.623 (0.0419) | [0.536, 0.700] | 0.0810 (0.160) | [-0.230, 0.392] | 0.000210 (0.000450) | [$1.22\times10^{-5}$, 0.00122] |
| Combinatorics | 1 | Y | 0.376 (0.0192) | [0.339, 0.414] | 0.000210 (0.162) | [-0.342, 0.281] | 0.00204 (0.00152) | [0.000229, 0.00601] |
| Comb. + length | 1 | Y | 0.378 (0.0190) | [0.341, 0.415] | 0.0801 (0.165) | [-0.272, 0.368] | 0.0219 (0.00812) | [0.00972, 0.0409] |
| Stability | 1 | Y | 0.675 (0.0253) | [0.624, 0.723] | 0.0529 (0.170) | [-0.306, 0.352] | $1.37\times10^{-5}$ ($9.96\times10^{-6}$) | [$5.25\times10^{-6}$, $3.64\times10^{-5}$] |
| Stab. + length | 1 | Y | 0.679 (0.0250) | [0.629, 0.727] | 0.122 (0.167) | [-0.237, 0.416] | $3.29\times10^{-5}$ ($2.73\times10^{-5}$) | [$7.40\times10^{-6}$, $8.98\times10^{-5}$] |
| Combinatorics | 2 | Y | 0.466 (0.0330) | [0.400, 0.529] | 0.000394 (0.162) | [-0.315, 0.318] | 0.00186 (0.00157) | [0.000232, 0.00602] |
| Comb. + length | 2 | Y | 0.473 (0.0326) | [0.408, 0.535] | 0.0502 (0.167) | [-0.276, 0.374] | 0.00495 (0.00367) | [0.000926, 0.0146] |
| Stability | 2 | Y | 0.602 (0.0325) | [0.536, 0.663] | 0.0606 (0.163) | [-0.258, 0.377] | 0.000235 (0.000289) | [$2.69\times10^{-5}$, 0.000991] |
| Stab. + length | 2 | Y | 0.604 (0.0324) | [0.538, 0.665] | 0.103 (0.166) | [-0.223, 0.421] | 0.000609 (0.000701) | [$5.19\times10^{-5}$ ,0.00251] |
| Combinatorics | 3 | Y | 0.631 (0.0336) | [0.563, 0.694] | $-5.16\times10^{-5}$ (0.162) | [-0.306, 0.325] | $4.44\times10^{-5}$ ($6.24\times10^{-5}$) | [$6.14\times10^{-6}$, 0.000210] |
| Comb. + length | 3 | Y | 0.643 (0.0319) | [0.578, 0.703] | 0.0409 (0.167) | [-0.276, 0.373] | $9.01\times10^{-5}$ (0.000107) | [$1.03\times10^{-5}$, 0.000359] |
| Stability | 3 | Y | 0.731 (0.300) | [0.669, 0.786] | 0.0197 (0.167) | [-0.297, 0.350] | $1.09\times10^{-5}$ ($6.19\times10^{-6}$) | [$5.15\times10^{-6}$, $2.68\times10^{-5}$] |
| Stab. + length | 3 | Y | 0.734 (0.0290) | [0.674, 0.788] | 0.0581 (0.171) | [-0.268, 0.397] | $1.28\times10^{-5}$ ($1.15\times10^{-5}$) | [$5.19\times10^{-6}$, $3.61\times10^{-5}$] |
| Combinatorics | 4 | Y | 0.624 (0.0355) | [0.551, 0.690] | $-7.38\times10^{-5}$ (0.163) | [-0.309, 0.323] | $5.15\times10^{-5}$ ($7.59\times10^{-5}$) | [$6.07\times10^{-6}$, 0.000238] |
| Comb. + length | 4 | Y | 0.666 (0.0305) | [0.603, 0.722] | 0.0367 (0.166) | [-0.277, 0.366] | $4.46\times10^{-5}$ ($5.56\times10^{-5}$) | [$5.36\times10^{-6}$, 0.000198] |
| Stability | 4 | Y | 0.692 (0.0389) | [0.609, 0.761] | 0.0237 (0.165) | [-0.290, 0.353] | $3.83\times10^{-5}$ ($5.52\times10^{-5}$) | [$5.89\times10^{-6}$, 0.000154] |
| Stab. + length | 4 | Y | 0.715 (0.0371) | [0.636, 0.780] | 0.0677 (0.166) | [-0.250, 0.396] | $2.87\times10^{-5}$ ($4.11\times10^{-5}$) | [$6.57\times10^{-6}$, 0.000122] |

† Yes (Y)/No (N) indicates exclusion/inclusion of sequences 'ACACACACAC' and 'CACACACACACACA', respectively.

**Supplementary Table 3:** Statistics of optimised parameters for various applications of the model on experimental data.

| Model | n | Outliers excluded† | $\kappa$ ($\times 10^6$ $s^{-1}$) mean (std-dev) | 95% CI | $\alpha$ mean (std-dev) | 95% CI | $\gamma$ mean (std-dev) | 95% CI |
|---|---|---|---|---|---|---|---|---|
| Combinatorics | 1 | N | 6.12 (0.103) | [5.92, 6.32] | -2‡ (-) | - | $-\infty$‡ (-) | - |
| Comb. + length | 1 | N | 5.12 (1.29) | [3.06, 8.07] | -1.92 (0.0997) | [-2.11, -1.72] | $-\infty$‡ (-) | - |
| Stability | 1 | N | 28.4 (5.35) | [20.6, 41.3] | -2‡ (-) | - | 6.56 (0.320) | [5.96, 7.22] |
| Stab. + length | 1 | N | 33.2 (9.18) | [18.8, 54.4] | -2.06 (0.105) | [-2.26, -1.85] | 6.55 (0.325) | [5.95, 7.22] |
| Combinatorics | 2 | N | 10.3 (0.178) | [10.0, 10.7] | -2‡ (-) | - | $-\infty$‡ (-) | - |
| Comb. + length | 2 | N | 6.63 (1.49) | [4.19, 10.0] | -1.80 (0.0936) | [-1.99, -1.62] | $-\infty$‡ (-) | - |
| Stability | 2 | N | 26.2 (5.38) | [13.0, 35.0] | -2‡ (-) | - | 7.50 (0.753) | [5.15, 8.29] |
| Stab. + length | 2 | N | 25.1 (11.2) | [7.68, 50.2] | -1.96 (0.117) | [-2.18, -1.73] | 7.40 (0.867) | [5.08, 8.32] |
| Combinatorics | 3 | N | 13.6 (0.243) | [13.1, 14.1] | -2‡ (-) | - | $-\infty$‡ (-) | - |
| Comb. + length | 3 | N | 8.01 (1.60) | [5.34, 11.6] | -1.76 (0.0876) | [-1.93, -1.59] | $-\infty$‡ (-) | - |
| Stability | 3 | N | 19.4 (0.653) | [18.1, 20.7] | -2‡ (-) | - | 8.17 (0.143) | [7.87, 8.44] |
| Stab. + length | 3 | N | 16.3 (3.49) | [10.5, 24.1] | -1.92 (0.0895) | [-2.09, -1.74] | 8.17 (0.146) | [7.86, 8.43] |
| Combinatorics | 4 | N | 14.7 (0.271) | [14.2, 15.2] | -2‡ (-) | - | $-\infty$‡ (-) | - |
| Comb. + length | 4 | N | 6.73 (1.19) | [4.70, 9.34] | -1.63 (0.0821) | [-1.79, -1.47] | $-\infty$‡ (-) | - |
| Stability | 4 | N | 46.4 (5.33) | [36.6, 56.7] | -2‡ (-) | - | 13.2 (0.499) | [12.7, 13.6] |
| Stab. + length | 4 | N | 27.6 (6.77) | [16.1, 42.0] | -1.74 (0.0881) | [-1.91, -1.56] | 13.2 (1.10) | [12.7, 13.8] |
| Combinatorics | 1 | Y | 5.99 (0.104) | [5.79,6.19] | -2‡ (-) | - | $-\infty$‡ (-) | - |
| Comb. + length | 1 | Y | 5.57 (1.45) | [3.25,8.90] | -1.96 (0.103) | [-2.16, -1.76] | $-\infty$‡ (-) | - |
| Stability | 1 | Y | 23.0 (3.41) | [17.8,31.1] | -2‡ (-) | - | 6.18 (0.283) | [5.65, 6.77] |
| Stab. + length | 1 | Y | 30.7 (9.07) | [16.8,51.9] | -2.11 (0.106) | [-2.31, -1.90] | 6.16 (0.282) | [5.65, 6.75] |
| Combinatorics | 2 | Y | 10.6 (0.186) | [10.3,11.0] | -2‡ (-) | - | $-\infty$‡ (-) | - |
| Comb. + length | 2 | Y | 7.79 (1.79) | [4.85, 11.8] | -1.86 (0.0948) | [-2.04, -1.67] | $-\infty$‡ (-) | - |
| Stability | 2 | Y | 20.4 (2.92) | [15.5, 26.7] | -2‡ (-) | - | 6.76 (0.449) | [5.85, 7.57] |
| Stab. + length | 2 | Y | 19.6 (6.09) | [5.83, 7.58] | -1.97 (0.0988) | [-2.16, -1.78] | 6.75 (0.456) | [5.83, 7.58] |
| Combinatorics | 3 | Y | 15.2 (0.263) | [14.7, 15.7] | -2‡ (-) | - | $-\infty$‡ (-) | - |
| Comb. + length | 3 | Y | 9.55 (1.89) | [6.36, 13.8] | -1.79 (0.0852) | [-1.96,-1.62] | $-\infty$‡ (-) | - |
| Stability | 3 | Y | 20.1 (0.795) | [18.6, 21.8] | -2‡ (-) | - | 7.75 (0.211) | [7.32, 8.15] |
| Stab. + length | 3 | Y | 16.9 (3.61) | [10.9, 24.9] | -1.91 (0.0872) | [-2.08, -1.74] | 7.74 (0.217) | [7.29, 8.15] |
| Combinatorics | 4 | Y | 17.4 (0.303) | [16.8, 18.0] | -2‡ (-) | - | $-\infty$‡ (-) | - |
| Comb. + length | 4 | Y | 6.92 (1.18) | [4.88, 9.51] | -1.57 (0.0767) | [-1.72, -1.42] | $-\infty$‡ (-) | - |
| Stability | 4 | Y | 22.2 (2.72) | [18.4, 29.3] | -2‡ (-) | - | 9.96 (0.772) | [8.32, 11.4] |
| Stab. + length | 4 | Y | 10.4 (2.62) | [6.15, 16.2] | -1.62 (0.0826) | [-1.78, -1.46] | 9.95 (1.35) | [7.28, 12.0] |

† Yes (Y)/No (N) indicates exclusion/inclusion of sequences 'ACACACACAC' and 'CACACACACACACA', respectively.

‡ These coefficients are fixed and are not allowed to freely vary. In the case of $\gamma$, the value should be considered in the appropriate limit such that all complementary binding states possess a probability of hybridisation of 1 and all mis-matched sites possess a probability of 0.

**Supplementary Table 4:** Point estimates for $\rho$, $R^2$, and model parameters using mean experimental rates as point estimates.

| Model | n | Outliers excluded† | $\rho$ | $R^2$ | $\kappa$ ($\times 10^6$ $s^{-1}$) | $\alpha$ | $\gamma$ |
|---|---|---|---|---|---|---|---|
| Combinatorics | 1 | N | 0.381 | 0.122 | 6.12 | -2‡ | $-\infty$‡ |
| Comb. + length | 1 | N | 0.383 | 0.124 | 4.97 | -1.92 | $-\infty$‡ |
| Stability | 1 | N | 0.712 | 0.502 | 27.5 | -2‡ | 6.55 |
| Stab. + length | 1 | N | 0.712 | 0.503 | 31.5 | -2.06 | 6.54 |
| Combinatorics | 2 | N | 0.514 | 0.249 | 10.3 | -2‡ | $-\infty$‡ |
| Comb. + length | 2 | N | 0.521 | 0.259 | 6.46 | -1.80 | $-\infty$‡ |
| Stability | 2 | N | 0.622 | 0.340 | 26.9 | -2‡ | 7.70 |
| Stab. + length | 2 | N | 0.622 | 0.340 | 25.0 | -1.97 | 7.68 |
| Combinatorics | 3 | N | 0.580 | 0.163 | 13.6 | -2‡ | $-\infty$‡ |
| Comb. + length | 3 | N | 0.589 | 0.180 | 7.85 | -1.76 | $-\infty$‡ |
| Stability | 3 | N | 0.679 | 0.309 | 19.4 | -2‡ | 8.18 |
| Stab. + length | 3 | N | 0.681 | 0.312 | 15.9 | -1.91 | 8.18 |
| Combinatorics | 4 | N | 0.564 | -0.0878 | 14.7 | -2‡ | $-\infty$‡ |
| Comb. + length | 4 | N | 0.583 | -0.0397 | 6.63 | -1.63 | $-\infty$‡ |
| Stability | 4 | N | 0.638 | -0.153 | 46.3 | -2‡ | 13.2 |
| Stab. + length | 4 | N | 0.644 | -0.130 | 27.3 | -1.74 | 13.3 |
| Combinatorics | 1 | Y | 0.389 | 0.127 | 59.9 | -2‡ | $-\infty$‡ |
| Comb. + length | 1 | Y | 0.390 | 0.128 | 5.40 | -1.96 | $-\infty$‡ |
| Stability | 1 | Y | 0.699 | 0.478 | 22.4 | -2‡ | 6.16 |
| Stab. + length | 1 | Y | 0.701 | 0.481 | 29.2 | -2.11 | 6.15 |
| Combinatorics | 2 | Y | 0.484 | 0.234 | 10.6 | -2‡ | $-\infty$‡ |
| Comb. + length | 2 | Y | 0.489 | 0.239 | 7.60 | -1.86 | $-\infty$‡ |
| Stability | 2 | Y | 0.623 | 0.384 | 20.1 | -2‡ | 6.78 |
| Stab. + length | 2 | Y | 0.622 | 0.384 | 18.6 | -1.97 | 6.76 |
| Combinatorics | 3 | Y | 0.655 | 0.429 | 15.2 | -2‡ | $-\infty$‡ |
| Comb. + length | 3 | Y | 0.665 | 0.443 | 9.36 | -1.79 | $-\infty$‡ |
| Stability | 3 | Y | 0.758 | 0.574 | 20.1 | -2‡ | 7.76 |
| Stab. + length | 3 | Y | 0.759 | 0.576 | 16.5 | -1.91 | 7.75 |
| Combinatorics | 4 | Y | 0.647 | 0.400 | 17.4 | -2‡ | $-\infty$‡ |
| Comb. + length | 4 | Y | 0.689 | 0.470 | 68.3 | -1.57 | $-\infty$‡ |
| Stability | 4 | Y | 0.717 | 0.458 | 21.6 | -2‡ | 9.98 |
| Stab. + length | 4 | Y | 0.738 | 0.507 | 9.95 | -1.64 | 9.99 |

† Yes (Y)/No (N) indicates exclusion/inclusion of sequences 'ACACACACAC' and 'CACACACACACACA', respectively.

‡ These coefficients are fixed and are not allowed to freely vary. In the case of $\gamma$, the value should be considered in the appropriate limit such that all complementary binding states possess a probability of hybridisation of 1 and all mis-matched sites possess a probability of 0.
