## Supplementary Note for "Mechanisms underlying sequence-dependent DNA hybridisation rates in the absence of secondary structure"

### Supplementary Note 1: Effective rates/mean first passage times for a three state system

Consider the three state system with constant transition rates

$$\mathrm{A} \underset{\mathrm{k_2}}{\overset{\mathrm{k_1}}{\rightleftharpoons}} \mathrm{B} \xrightarrow{\mathrm{k_3}} \mathrm{C},$$

with the rate for transitions $C \rightarrow B$ either naturally or artifically supressed such that state $C$ acts as an absorbing state. Here we will calculate the effective rate of transitioning from state A to state C, $k^{\mathrm{eff}}_{A\rightarrow C}$, characterised by the inverse of the mean first passage time (MFPT), $\langle t^{\mathrm{FP}}_{A\rightarrow C}\rangle$.

The distribution of waiting times to transition out of any state $x$ is $\lambda_x e^{-\lambda_x t}$, by assumption, where $\lambda_x = \sum_{y\neq x} k_{x\rightarrow y}$, such that the expected time from arrival into state $x$ before a transition is $t_x = 1/\lambda_x$. From a given state $x$ the probability that such a transition is into some specific state $z$ is $P_{x\rightarrow z} = k_{x\rightarrow z}/\lambda_x$. Given a sequence of transitions from state $x$, the expected number of transitions required to first observe the specific transition $x \rightarrow z$ is the expectation $\langle n_{x\rightarrow z}\rangle = \sum_{i=1}^{\infty} iP_{x\rightarrow z}(1 - P_{x\rightarrow z})^{i-1} = 1/P_{x\rightarrow z}$.

Given the transition network between states $A$, $B$, and $C$, we can consequently expect $\langle n_{B\rightarrow C}\rangle = 1/P_{B\rightarrow C} = (k_2 + k_3)/k_2$ transitions from state $B$ consisting of $\langle n_{B\rightarrow C}\rangle - 1$ transitions $B \rightarrow A$ followed by the final transition $B \rightarrow C$. Each of these is paired with a preceding $A \rightarrow B$ transition. Consequently the mean time to reach state $C$ is given by

$$\begin{aligned} t^{\mathrm{FP}}_{A\rightarrow C} &= \langle n_{B\rightarrow C}\rangle \left(t_A + t_B\right) \\ &= \frac{k_2 + k_3}{k_3}\left(\frac{1}{\lambda_A} + \frac{1}{\lambda_B}\right) \\ &= \frac{k_2 + k_3}{k_3}\left(\frac{1}{k_1} + \frac{1}{k_2 + k_3}\right) \\ &= \frac{k_1 + k_2 + k_3}{k_1 k_3}. \end{aligned} \tag{1}$$

We then identify the rate as the inverse of this quantity viz.

$$k^{\mathrm{eff}}_{A\rightarrow C} = \frac{1}{\langle t^{\mathrm{FP}}_{A\rightarrow C}\rangle} = \frac{k_1 k_3}{k_1 + k_2 + k_3}. \tag{2}$$

### Supplementary Note 2: Statistical Approaches

Here we will cover the approaches taken for generating estimates and confidence intervals for reported correlations, $R^2$ values, and, $p$-values. We shall also provide definitions of these quantities where appropriate and point to how they might be used to infer significance and over-fitting.

#### Definitions

We will be assuming the existence of indexed data of size $N$, $d = \{d_i \mid i \in \{1 \ldots N\}\}$, consisting of 3-tuples $d_i = \{s_i, x_i, \varepsilon_i\}$, where $s_i$ are individual $5' - 3'$ DNA sequences forming indexed set $s = \{s_i \mid i \in \{1 \ldots N\}\}$, $x_i$ are (mean) experimental hybridisation rates forming indexed set $x = \{x_i \mid i \in \{1 \ldots N\}\}$, and $\varepsilon_i$ are standard deviations forming indexed set $\varepsilon = \{\varepsilon_i \mid i \in \{1 \ldots N\}\}$.

The hybridisation model then generates rates $y = \{y_i \mid i \in \{1 \ldots N\}\}$, where we can write $y = \mathcal{M}(x, s)$ to emphasise that the $y_i$ values are a function, $\mathcal{M}$, of both the sequences $s$ *and* the experimental hybridisation rates $x$. Explicitly, this function subsumes all optimisation of internal parameters of the model to best generate an output $y$ which is similar to the provided $x$. We then report goodness of fit measures between $x$ and the resultant $y$. Specifically we use the Pearson correlation coefficient

$$\rho(x, y) = \frac{\sum_{i=1}^{N}(x_i - \bar{x})(y_i - \bar{y})}{\sqrt{\sum_{i=1}^{N}(x_i - \bar{x})^2 \cdot \sum_{i=1}^{N}(x_i - \bar{x})^2}},$$

and coefficient of determination, or $R^2$,

$$R^2(x, y) = 1 - \frac{\sum_{i=1}^{N}(y_i - x_i)^2}{\sum_{i=1}^{N}(y_i - \bar{y})^2},$$

where $\bar{x}$ and $\bar{y}$ are the means of $x$ and $y$, respectively. Note, the nature of the model (being absent a freely variable intercept parameter), means that generally $R^2 \neq \rho^2$, and that there is no requirement for $R^2 \geq 0$.

#### Point estimates

Goodness of fit measures, model hybridisation rates and parameters for point estimates of the experimental hybridisation rates are based on the values $x$ and the subsequent values $y = \mathcal{M}(x, s)$ with no reference to $\varepsilon$.

#### Error analysis of estimates

We may provide estimates of the statistics of the model (e.g. means, standard deviations and confidence intervals of goodness-of-fit measures, parameters and rates) by recognising that the provided data $x$ are in fact estimates of the (unknown) true values. To find statistics of our measures we re-sample our original data under the assumption that the errors are normally distributed with standard deviation equal to the provided experimental standard deviation of the individual experimental obervations, $\varepsilon$.

As such we generate $n_s$ re-sampled data sets, $\hat{x}^{(i)}$, $i \in \{1 \ldots n_s\}$ according to

$$\hat{x}^{(i)} = \left\{ \hat{x}_j^{(i)} = x_j + z_j^{(i)} \mid z_j^{(i)} \sim \mathcal{N}(0, \varepsilon_j),\ j \in \{1 \ldots N\} \right\}$$

where $z \sim \mathcal{N}(0, \varepsilon_j)$ indicates that $z$ is a normally distributed random number with mean 0 and standard deviation $\varepsilon_j$.

For each generated data set, $\hat{x}^{(i)}$, we run the model to get a set of estimates, $\hat{y}^{(i)} = \mathcal{M}(\hat{x}^{(i)}, s)$, from which we obtain an estimate of the true correlation and coefficient of determination, $\hat{\rho}_{(i)} = \rho(\hat{x}^{(i)}, \hat{y}^{(i)})$ and $\hat{R}^2_{(i)} = R^2(\hat{x}^{(i)}, \hat{y}^{(i)})$, in turn leading to a distribution of $\hat{\rho} = \{\hat{\rho}_{(i)} \mid i \in \{1 \ldots n_s\}\}$, and $\hat{R}^2 = \{\hat{R}^2_{(i)} \mid i \in \{1 \ldots n_s\}\}$ from which we can report estimates of the mean, median, standard deviation, and confidence intervals. This can then also be done analogously for model parameters and hybridisation rates.

#### Determining $p$-values and assessing over-fitting with permutation tests

We provide a measure of statistical significance by calculating standard right-tailed $p$-values for the calculations of the goodness-of-fit statistics $\rho$ and $R^2$. As per usual, smaller values indicate that a result equally or more extreme (values of $\rho$ or $R^2$ greater or equal to those for the model and experimental data), is less likely under the null distribution.

To determine these $p$-values we must be able to specify, or sample from, such a null distribution for our test statistics. Direct specification is not available here, and so a standard way of sampling such a distribution is through repeated generation of random permutations of the provided data set, and calculating the test statistic for each such permutation. The implicit assumption is that data is exchangeable under the null hypothesis. For us this means the null hypothesis is that there is no relationship between i) DNA strand sequences or lengths and ii) their hybridisation rates. Note, for instance, that under the null hypothesis we would not expect similar strand sequences to possess similar hybridisation rates.

The samples from that null distribution can then also serve as suitably random data to assess how well the model can fit to arbitrary patterns, thus providing a rudimentary insight into the possibility of over-fitting.

##### Sampling from the null distribution

For our purposes a permutation is understood as a function which 'shuffles' the data-values (the hybridisation rates, $x_i$), but not the labels (the sequences, $s_i$). A given permutation, or shuffling, indexed by $i$, can be represented by notation $\sigma_i$. This can be understood as a bijective function on data indices $k$, $\sigma_i(k) : \{1 \dots N\} \to \{1 \dots N\}$. As such given data indices $\{1, 2, 3, 4, 5\}$, a permutation $\sigma_i$ might be $\{2, 5, 1, 3, 4\}$, such that $\sigma_i(5) = 4$ and $\sigma_i(3) = 1$. As such, a single element, $d_k$, of our data $d$ under permutation $\sigma_i$ can be written

$$d_{\sigma_i(k)} = \{s_k, x_{\sigma_i(k)}, \varepsilon_{\sigma_i(k)}\}$$

leading to permuted sets $x_{\sigma_i} = \{x_{\sigma_i(k)} \mid k \in \{1 \dots N\}\}$, and $\varepsilon_{\sigma_i} = \{\varepsilon_{\sigma_i(k)} \mid k \in \{1 \dots N\}\}$. To account for experimental uncertainty, we simlutaneously resample from the assumed normally distributed standard deviations to give a set of samples from the null distribution

$$\tilde{x}_{\sigma_i} = \left\{ \tilde{x}_{\sigma_i(k)} = x_{\sigma_i(k)} + z_k^{(i)} \mid z_k^{(i)} \sim \mathcal{N}(0, \varepsilon_{\sigma_i(k)}),\ k \in \{1 \dots N\} \right\}.$$

Using these sets we can find model predictions for the samples from the null distribution viz.

$$\tilde{y}^{(i)} = \mathcal{M}(\tilde{x}_{\sigma_i}, s).$$

This in turn gives estimates of correlation and coefficients of determination for the samples

$$\begin{aligned} \tilde{\rho}_{(i)} &= \rho(\tilde{x}_{\sigma_i}, \tilde{y}^{(i)}), \\ \tilde{R}^2_{(i)} &= R^2(\tilde{x}_{\sigma_i}, \tilde{y}^{(i)}), \end{aligned}$$

thus yielding null-distributed sets $\tilde{\rho} = \{\tilde{\rho}_{(i)} \mid i \in \{1 \dots n_p\}\}$, and $\tilde{R}^2 = \{\tilde{R}^2_{(i)} \mid i \in \{1 \dots n_p\}\}$, where $n_p$ is the number of such random permutations.

##### Use of null distributed statistics for over-fitting detection

It is important to recognise that our test statistics are the goodness-of-fit measures of the model with the data after optimisation of the internal parameters of the model to best fit the provided data. As such running the model on each permutation $\tilde{x}_{\sigma_i}$, indicated here through function $\mathcal{M}(\tilde{x}_{\sigma_i}, s)$, independently attempts to find the best fit under that particular permutation with a separately optimised set of internal model parameters. As such the null distributed values $\tilde{\rho}$ and $\tilde{R}^2$ can be used to infer over-fitting through the model's ability to replicate/correlate with random data[1]. For instance if the mean of $\tilde{\rho}$ is $\sim 0$, the model is broadly unable to fit arbitrarily varying data, whereas if the mean of $\tilde{\rho}$ is high it indicates the model has enough free internal structure to fit to any data presented to it. Such properties cannot be determined from $p$-values alone and so an 'effect size' can also be considered consisting of the difference between mean goodness-of-fit statistics ($\rho(\cdot, \cdot)$, $R^2(\cdot, \cdot)$) resulting from the data and resulting from the null distribution. Explicitly, if a high correlation, e.g. $\rho = 0.9$, is found with data, but the mean of $\tilde{\rho}$ is also high, e.g. $\langle \tilde{\rho} \rangle = 0.8$, then even if the result is significant, much of the correlation may be arising from excess model complexity.

[1] distributed under the null distribution.

##### $p$-value estimates for point-estimates of $\rho$ and $R^2$

Since $\tilde{\rho}$ and $\tilde{R^2}$ are sampled from the null distribution, we can estimate a $p$-value for singly specified values of $\rho$ and $R^2$ by determining the proportion of $\tilde{\rho}$ and $\tilde{R^2}$ greater than $\rho$ and $R^2$, respectively. Since the number of generated permutations in practice is less than all possible permutations[2] any such proportion should be considered to be an estimate, $\hat{p}$, of the true $p$-value, $p$. An un-biased estimator of $p$ is given by

$$\hat{p} = \frac{g}{n_p}$$

where $g$ is the number of permutations with $\tilde{\rho}_{(i)}$ or $\tilde{R}^2_{(i)}$ greater than the specified $\rho$ or $R^2$, respectively, and as before, $n_p$ the number of permutations. However, this can lead to failures to protect against type-I errors at the specifed confidence level [1]. This can be avoided through the use of the conservatively biased estimator [1]

$$\hat{p}_{\text{bias}} = \frac{g+1}{n_p+1}.$$

The number of permutations with a larger value of the relevant statistic, $g$, is effectively drawn from a binomial distribution. As such the standard deviation associated with estimates of $p$ (equivalent to the standard error for the Bernoulli variable of whether a single permutation's statistic is more extreme than the data) can be specified as

$$\text{std-dev}_p = \sqrt{\frac{\hat{p}_{\text{bias}}(1-\hat{p}_{\text{bias}})}{n_p}} + \mathcal{O}(n_p^{-\frac{3}{2}}).$$

In turn, confidence intervals for the estimate of $p$ can be specified through standard library functions. Specifically we report the "exact" Clopper-Pearson intervals at various confidence levels in the relevant data files.

##### $p$-value estimates for distributions of estimates of $\rho$ and $R^2$

In the case where distributions of $n_s$ estimates $\hat{\rho}$ and $\hat{R}^2$ are specified, we must account for the two sources of variability, namely that arising from the variation across the $n_s$ values of $\hat{\rho}_{(i)}$ and $\hat{R}^2_{(i)}$, and the fact that any single estimated $p$-value from these distibutions is itself a point estimate, with associated variance, confidence etc.

To achieve this, for each estimate $\hat{\rho}_{(i)}$ or $\hat{R}^2_{(i)}$ $(i \in \{1, n_s\})$, we first calculate the number of permutations, $g_{(i)}$, for which the permuted values $\tilde{\rho}_{(j)}$ and $\tilde{R^2}_{(j)}$ $(j \in \{1 \ldots n_p\})$ are greater than those estimates as per the previous section. This leads to unbiased estimates of the $p$-value $\hat{p}_{(i)} = g_{(i)}/n_p$. We then exploit the property that such an estimate arises from a binomial distribution associated with the permutation procedure. As such the probability of getting exactly $g$ such cases can be written

$$P_{\text{binom}}(g; n_p, p) = \binom{n_p}{g} p^g (1-p)^{n-g}.$$

This is then reinterpreted as a likelihood function for the true value $p$

$$\mathcal{L}_{\text{binom}}(p; g, n_p) = \binom{n_p}{g} p^g (1-p)^{n-g},$$

for which the estimate $\hat{p} = g/n_p$ is the maximum likelihood estimate (MLE) $\hat{p} = \arg\max_p L(p; g, n_p)$. We may convert this to a probability density viz.

$$\mathcal{P}(p; g, n_p) = \frac{\mathcal{L}_{\text{binom}}(p; g, n_p)\mathcal{P}_{\text{null}}(p)}{\int_0^1 dp\, \mathcal{L}_{\text{binom}}(p; g, n_p)\mathcal{P}_{\text{null}}(p)},$$

where $\mathcal{P}_{\text{null}}(p)$ is the prior distribution on $p$ under the null, which is by definition uniform [1], such that

$$\begin{aligned}
\mathcal{P}(p; g, n_p) &= \frac{\mathcal{L}_{\text{binom}}(p; g, n_p)}{\int_0^1 dp\, \mathcal{L}_{\text{binom}}(p; g, n_p)} \\
&= \frac{p^g (1-p)^{n_p-g}}{\int_0^1 dp\, \hat{p}^g (1-p)^{n_p-g}} \\
&= \frac{\Gamma(n_p+2)}{\Gamma(n_p-g+1)\Gamma(g+1)} p^g (1-p)^{n_p-g} \\
&= \beta(p; g+1, n_p-g+1),
\end{aligned}$$

[2] The total number of permutations is $N!$. E.g. for $N = 40$ there are approximately $8.16 \times 10^{47}$ possible permutations.

and where $\Gamma(\cdot)$ & $\beta(\cdot;\cdot,\cdot)$ are the gamma function and beta distribution, respectively.

Consequently, for each value $\hat{p}_{(i)} = g_{(i)}/n_p$, for a given estimate $\hat{\rho}_{(i)}$ or $\hat{R}^2_{(i)}$, we account for the fact that $\hat{p}_{(i)}$ is merely a point estimate by generating some number, $n_b$, beta distributed $p$-values drawn from $\beta(g_{(i)}/n_p; g_{(i)} + 1, n_p - g_{(i)} + 1)$. We then convert to the conservative biased estimate, *c.f.* the previous section, such that we have a sub-ensemble

$$\tilde{p}_{(i)} = \left\{ \tilde{p}_{(i),j} = \frac{z_j^{(i)} n_p + 1}{n_p + 1} \mid z_j^{(i)} \sim \beta(\hat{p}_{(i)}; g_{(i)} + 1, n_p - g_{(i)} + 1),\ j \in \{1 \ldots n_b\} \right\}.$$

We then combine all such sub-ensembles to create a total distribution of possible $p$-values

$$\tilde{p} = \left\{ \tilde{p}_{(i),j} \mid \tilde{p}_{(i),j} \in \tilde{p}_{(i)},\ i \in \{1 \ldots n_s\},\ j \in \{1 \ldots n_b\} \right\},$$

from which we may compute mean values, standard deviations and confidence intervals.
